# COALIA: a computational model of human EEG for consciousness research

**DOI:** 10.1101/575043

**Authors:** Siouar Bensaid, Julien Modolo, Isabelle Merlet, Fabrice Wendling, Pascal Benquet

**Author notes:** Co-first authors (Siouar Bensaid, Julien Modolo). Correspondence: Fabrice Wendling.

## Abstract

Understanding the origin of the main physiological processes involved in consciousness is a major challenge of contemporary neuroscience, with crucial implications for the study of Disorders of Consciousness (DOC). The difficulties in achieving this task include the considerable quantity of experimental data in this field, along with the non-intuitive, nonlinear nature of neuronal dynamics. One possibility of integrating the main results from the experimental literature into a cohesive framework, while accounting for nonlinear brain dynamics, is the use of physiologically-inspired computational models. In this study, we present a physiologically-grounded computational model, attempting to account for the main micro-circuits identified in the human cortex, while including the specificities of each neuronal type. More specifically, the model accounts for thalamo-cortical (vertical) regulation of cortico-cortical (horizontal) connectivity, which is a central mechanism for brain information integration and processing. The distinct neuronal assemblies communicate through feedforward and feedback excitatory and inhibitory synaptic connections implemented in a template brain accounting for long-range connectome. The EEG generated by this physiologically-based simulated brain is validated through comparison with brain rhythms recorded in humans in two states of consciousness (wakefulness, sleep). Using the model, it is possible to reproduce the local disynaptic disinhibition of basket cells (fast GABAergic inhibition) and glutamatergic pyramidal neurons through long-range activation of VIP interneurons that induced inhibition of SST interneurons. The model (COALIA) predicts that the strength and dynamics of the thalamic output on the cortex control the local and long-range cortical processing of information. Furthermore, the model reproduces and explains clinical results regarding the complexity of transcranial magnetic stimulation TMS-evoked EEG responses in DOC patients and healthy volunteers, through a modulation of thalamo-cortical connectivity that governs the level of cortico-cortical communication. This new model provides a quantitative framework to accelerate the study of the physiological mechanisms involved in the emergence, maintenance and disruption (sleep, anesthesia, DOC) of consciousness.

## 1 Introduction

The characterization and understanding of the mechanisms underlying consciousness is one, if not the greatest challenge that contemporary neuroscience is facing. Beyond the purely fundamental interest of this question, a major clinical issue is also at stake: evaluating residual consciousness in patients suffering from Disorders of Consciousness (DOC), which can be extremely difficult, and have crucial implications in terms of clinical care. For example, motor imagery paradigms can reveal covert consciousness in coma patients, using functional magnetic resonance imaging (fMRI) (Owen, Coleman et al. 2006) for instance. This illustrates the pressing need for an improved characterization of the mechanisms that underlie consciousness, which could be exploited to propose novel quantified measures, or *metrics*, of consciousness.

Many theories have attempted, at various levels of description, to integrate the multifaceted aspects of consciousness. One of the first theories that found a significant echo in the neuroscience community is the Dynamic Core Hypothesis (Tononi and Edelman 1998), which was the first to relate the concept of information with consciousness. In this theory, functional clusters in the thalamocortical system are central, and involve fast re-entrant interactions, as well as a high level of integration and differentiation giving rise to complex patterns of neuronal activity. Another popular theory that has been gradually expanded over the years, and that has solid ties with neurophysiology, is the Global Workspace Theory (GWT) (Dehaene, Kerszberg et al. 1998, Dehaene and Changeux 2011). In short, GWT states that conscious information is globally available within the brain, and that the “ignition” of large-scale networks, i.e. the sudden communication between distant brain regions to engage into the processing of information, enables a stimulus to reach the global workspace, hence consciousness. Ignition is thought to involve long-range glutamatergic fibers that enable long-distance communication between cortical regions. Several experiments have supported GWT, for example that non-masked words involve the activation of much wider networks as compared to masked words (Dehaene, Naccache et al. 2001), with a similar result found for sub-liminal versus supra-threshold visual stimuli (Modolo, Hassan et al. 2018, van Vugt, Dagnino et al. 2018). The Integrated Information Theory (IIT) is based on a different approach. Instead of beginning from the large-scale structure of the thalamo-cortical system as in the DCH and GWT, IIT introduces several axioms to derive general principles of consciousness. One of the leading ideas of IIT is that consciousness involves the *integration* of information between distant areas (reminiscent of ignition in GWT), which increases the complexity of the processed information. Segregation, or *differentiation* of information, is also key in IIT: for example, large-scale synchronization of several regions with the same activity is indeed integrated, however with low complexity (Koch, Massimini et al. 2016). Therefore, *integration* and *differentiation* appear as the two concepts leading to increasing the *complexity* of the information conveyed by brain-scale networks. A more recent theory named algorithmic information theory of consciousness a.k.a. Kolmogorov Theory (KT) (Ruffini 2017) is also based on the idea that conscious states are associated with higher levels of complexity, and that subjective experience occurs following processes of information compression.

In contrast with the aforementioned theoretical studies of consciousness, only few studies have actually attempted to simulate brain activity associated with consciousness states using neurophysiologically-plausible computational models. Obviously, the *in silico* implementation of neural mechanisms that underlie the emergence and maintenance of consciousness represents a considerable challenge. However, capturing the main features of the most significant common principles from the main theories of consciousness using a computational neuroscience framework appears at reach. For example, a computational model exploring how cortico-cortical connectivity is functionally impaired during sleep (“connectivity breakdown”) has been proposed (Esser, Hill et al. 2009). However, most of these models are limited in terms of spatial scale and the represented micro-circuitry. This limitation hinders bridging the micro-circuit scale with the brain-scale, which is of interest in consciousness. The present study proposes to fill this gap, and provides new links between different levels of description (from local neuronal population to whole-brain scale).

Using a bottom-up approach, we developed a new computational model of brain-scale electrophysiological activity. The model starts from neuronal micro-circuits involving different cellular subtypes that have been reliably identified through neurobiological studies. The basic unit of the model is the neural mass, representing a local population of a few thousands of neurons, which has proven its ability to capture the dynamics of actual neuronal assemblies (Wendling, Bartolomei et al. 2002). At the local level, the model includes subsets of pyramidal neurons (glutamatergic), and three different types of interneurons (GABAergic) with appropriate physiologically-based kinetics (fast *vs*. slow). At the global level, the large-scale model is then constructed on the basis of a standard 66-region brain atlas (Desikan, Segonne et al. 2006), with one neural mass representing the local field activity of one atlas region. Neural masses are spatially distributed over the cortex, using the template brain morphology (Colin). As they account for distinct cortical regions, neural masses are synaptically connected through long-range glutamatergic projections among pyramidal neurons and GABAergic interneurons, Connectivity is derived from DTI (Diffusion Tensor Imaging) data. Results show that the model captures the large-scale structure of brain connectivity between regions, while accounting for the properties of local micro-circuits. It can accurately reproduce EEG activity for different conscious states (e.g., sleep vs. wake), and the breakdown of functional connectivity during sleep as assessed through the replication of TMS-EEG experiments.

In this article, we first describe basic concepts of cortico-cortical and thalamo-cortical networks involved in theories of consciousness. Then, we present the neural mass modeling approach used to develop the brain-scale model, along with the various microcircuits considered and their functional role. A toy model involving a limited number of neuronal populations is then investigated in order to validate the implemented micro-circuits, before an extension to the whole-brain model. Simulated responses to TMS are generated and quantified in two consciousness states, namely awake and asleep. Results are discussed according to the novelty, performance and limitations of the model, along with its usefulness in consciousness studies. Future extensions are described.

## 2 Background: role of cortico- and thalamo-cortical networks in consciousness

Consciousness is a global functional state of the brain that is intrinsically linked with neuronal oscillations generated by large-scale cortico-cortical and thalamo-cortical networks (Llinas, Ribary et al. 1998). More specifically, wakefulness is determined by widespread thalamocortical projections (Timofeev and Steriade 1996, Laureys 2004) while awareness requires the activation of a wide cortico-cortical network, involving lateral and medial frontal regions parieto-temporal and posterior parietal areas, bilaterally (Laureys, Goldman et al. 1999). In the model, we included these two key components of consciousness that are briefly reviewed below.

### 2.1 “Horizontal” cortico-cortical connectivity

Functional connectivity studies have shed light on the functional networks involved in various conscious states (Jin and Chung 2012). During general anesthesia-induced loss of consciousness, there is a breakdown in cortical effective connectivity (Ferrarelli, Massimini et al. 2010, Hudetz 2012, Gomez, Phillips et al. 2013). As a reminder, effective connectivity is defined as the ability of a neuronal group to causally affect the firing of other neuronal groups (Friston 2011). In unresponsive patients, impaired consciousness was associated with altered effective connectivity (Varotto, Fazio et al. 2014, Crone, Schurz et al. 2015). A protocol of TMS triggered a simple local EEG response indicating a breakdown of effective connectivity at the cortical level, similar to the one previously observed in unconscious sleeping or anaesthetized subjects (Casali, Gosseries et al. 2013). Sleep stages have a drastic impact on consciousness and also on functional connectivity. For example, there is a strong reduction of both wakefulness and awareness components of consciousness during NREM sleep, associated with thalamic up-and-down state and cortical slow wave sleep. However, brain effective connectivity changes significantly (Tononi and Sporns 2003, Esser, Hill et al. 2009). More specifically, cortical activations become more local and stereotypical upon falling into NREM sleep, which indicates an impaired effective cortical connectivity (Massimini, Ferrarelli et al. 2010). “Horizontal” communication through coherence that involves high frequency oscillations synchronization is a fundamental mechanism in cortical function and perception (Fries 2005, Fries 2009). Overall, the “awareness” component of consciousness depends on large-scale synchronized communication among distant neuronal populations distributed over the neocortex (see Figure 1A, left panel).

**Figure 1.**
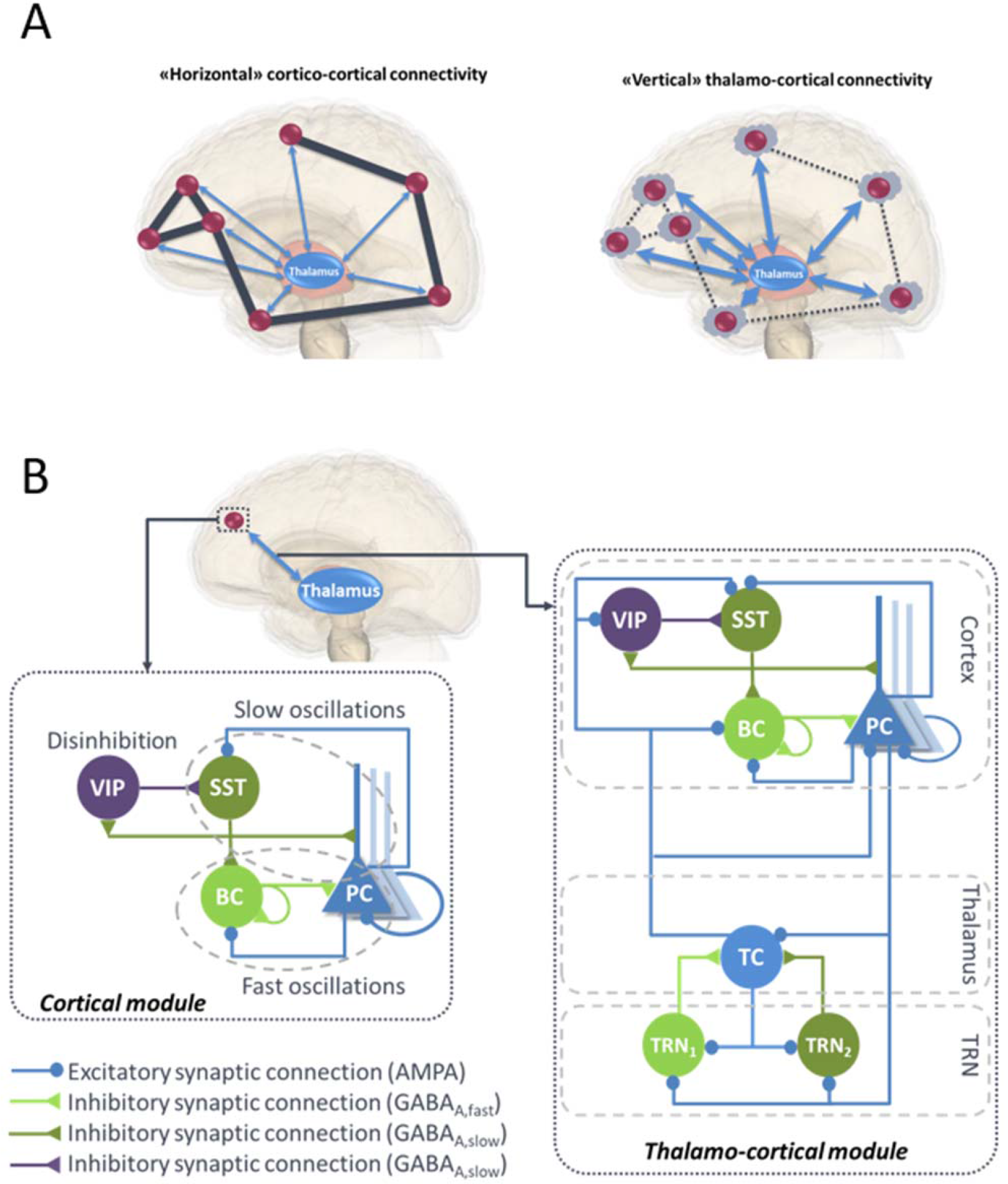
(A) Illustration of “horizontal” and “vertical” connectivity. ***Left:*** Horizontal connectivity refers to the cortico-cortical connections which are functionally effective during wakefulness, with a weak level of thalamo-cortical coupling. ***Right:*** Vertical connectivity refers to thalamo-cortical projections that functionally impair cortico-cortical connectivity during sleep. (**B) General architecture of the micro- and macro-circuits implemented in the computational model**. The local NMM of cortical activity is composed of a PC exciting two GABAergic interneurons, namely, the somatic-projecting BC and the dendritic-projecting SST, responsible for the generation of fast and slow oscillations, respectively; the VIP were introduced as they play a crucial role in cortical column communication through disinhibition of SST. The subcortical module consisted in TC sending excitatory glutamatergic projections to the TRN block composed of fast and slow GABAergic interneurons TRN_1_ and TRN_2_, respectively. **VIP:** vasoactive intestinal peptide positive GABAergic interneurons. **SST:** Somatostatin-positive GABAergic interneurons. **BC:** Basket-type GABAergic interneurons. **PC:** Glutamatergic Pyramidal Cells.

### 2.2 “Vertical” thalamo-cortical connectivity

As reported in IIT (Tononi 2004, Tononi 2012), consciousness depends on the brain’s ability to integrate information, which relies on the effective connectivity among functionally specialized regions (or clusters) of the thalamocortical system, and on the segregation of information. One important modulator of cortical connectivity is the activity pattern of thalamocortical cells, tonic *vs* up-and down, which is able to modify the excitability level of cortical neuron subpopulations. The thalamic-mediated synchronization of distant cortical areas may coordinate the large-scale integration of information across multiple cortical circuits, consequently influencing the level of arousal and consciousness (Saalmann 2014). Conversely, during sleep or anesthesia-induced transitions in consciousness, both thalamo-cortical and intra-thalamic functional connectivity are modified (Kim, Hwang et al. 2012, Hale, White et al. 2016). In addition, thalamic input to neocortex modifies cortico-cortical connectivity. Upon falling into NREM sleep (when rhythmic thalamo-cortical up-and-down activity develops), cortical activations become more local and stereotypical, indicating a significant decrease of cortico-cortical connectivity (Esser, Hill et al. 2009, Massimini, Ferrarelli et al. 2010, Usami, Matsumoto et al. 2015). Recently, it was shown that direct and tonic optogenetic activation of thalamic reticular nuclei (TRN) GABAergic interneurons induces a spatially restricted cortical slow wave activity (Lewis, Voigts et al. 2015). This activity was reminiscent of sleep rhythms, and animals exhibited behavioral changes that were consistent with a decrease of arousal.

Overall, this brief literature review suggests that both components of consciousness, namely awareness and wakefulness, are impaired when large-scale cortico-cortical functional connectivity mediated through the binding of synchronized high-frequency oscillations in the beta-gamma band (Schoenberg, Ruf et al. 2018) and regulated through the thalamus is (Nakajima and Halassa 2017). Meanwhile, during this decrease of awareness and wakefulness, an increase of “vertical” thalamo-cortical connectivity is observed, along with a stronger synchronization of delta oscillations between TC cell assemblies and isolated groups of neocortical neurons (Hill and Tononi 2005) (Figure 1A, right panel).

## 3 Materials and Methods

### 3.1 Modeling of micro- and macro-circuits: neural mass model approach

Neural mass models (NMMs) are a mathematical description of neural dynamics at a mesoscopic scale (from a millimeter to several centimeters of the cortex). This class of models was proposed in the 1970’s as an alternative to detailed microscopic models that require a more extensive computational cost (Wilson and Cowan 1973, Nunez 1974, Lopes da Silva, van Rotterdam et al. 1976, Freeman 1978). NMMs can indeed model the local field potential (LFP) of an entire cortical region using only few state variables (Breakspear 2017), whereas in detailed models this activity is meticulously described at the level of spatially distributed and interconnected neuron models, each including the properties of ionic channels, axons and dendrites (Wang and Buzsáki 1996, Whittington, Traub et al. 2000, Maex and De Schutter 2007).

Despite their simplicity, NMMs are neurophysiologically grounded, since they include the connectivity, synaptic kinetics and firing rates of neuronal sub-types present in the region of interest. The reduced complexity and performance (in term of reproducing actual LFPs) of the NMM approach made it a powerful tool to investigate various cerebral mechanisms, such as the generation of brain rhythms (Jansen and Rit 1995, David and Friston 2003, Ursino, Cona et al. 2010). NMMs have also been extensively used to study pathological dynamics such as in epilepsy (Wendling, Bartolomei et al. 2002, Traub, Contreras et al. 2005, Molaee-Ardekani, Benquet et al. 2010) (for a review, see (Wendling, Benquet et al. 2016)), Alzheimer’s disease (Bhattacharya, Coyle et al. 2011) and Parkinson’s disease (Liu, Wang et al. 2016, Liu, Zhu et al. 2017).

Designing a NMM involves identifying the main subsets of neurons implied in the modeled brain tissue and describing their synaptic interconnections. Based on an exhaustive literature review, we developed a models consisting of coupled NMMs able to simulate both cortical and thalamic activity, as described hereafter.

### 3.2 A local neural mass model of neocortical activity

Until recently, the classification of GABAergic interneurons was a highly challenging task, regarding the electrophysiological properties, morphology, biochemistry markers and connectivity. However, all recent studies converge towards much simpler functional categories, involving three main classes of GABAergic interneurons in the neocortex (Rudy, Fishell et al. 2011). As briefly reviewed below, these classes account for (1) somatic-targeting parvalbumine positive (PV^+^) basket cells (BC), (2) dendritic-targeting somatostatin positive (SST) interneurons and (3) vasoactive intestinal-peptide-(VIP) expressing interneurons (Tremblay, Lee et al. 2016), (Figure 1B, left panel).

#### 3.2.1 Basket cells and fast oscillations

PV^+^ participate to the generation of cortical gamma oscillations through: 1) thalamocortical feedforward inhibition in layer 4; 2) feedback inhibition in layer 2/3; and 3) via direct PV^+^/PV^+^ coupling through electrical gap-junctions (Povysheva, Zaitsev et al. 2008, Buzsáki and Wang 2012, Lewis, Curley et al. 2012, Varga, Oijala et al. 2014, Womelsdorf, Valiante et al. 2014, Chen, Zhang et al. 2017). Recent optogenetic experiments have demonstrated the causal role of somatic-targeting interneurons BC in mediating fast oscillations (> 20 Hz)(Chen, Zhang et al. 2017).

#### 3.2.2 Surround inhibition by SST GABAergic interneurons and slow oscillations

Neocortical SST neurons can exhibit high levels of spontaneous slow oscillations, and their tonic activity might facilitate fine scale up-and down regulation of global inhibition levels in the neocortex (Urban-Ciecko and Barth 2016). Lateral inhibition is a fundamental principle in neuronal networks (Harris and Mrsic-Flogel 2013, Karnani, Agetsuma et al. 2014, Harris and Gordon 2015). Lateral inhibition between nearby pyramidal cells (PCs) is thought to work through SST interneurons (Kapfer, Glickfeld et al. 2007, Silberberg and Markram 2007, Adesnik, Bruns et al. 2012). Dendritic inhibition is more effective than perisomatic inhibition in regulating excitatory synaptic integration, therefore SSTs are the key regulator of input-output transformations (Lovett-Barron, Turi et al. 2012). Furthermore, the slow kinetics of dendritic-targeting inhibitory postsynaptic membrane potentials (IPSPs) are particularly suited to maximize the localized shunting inhibition effect (Gidon and Segev 2012, Paulus and Rothwell 2016). Finally, it was shown that this cell type is especially involved in slow oscillations (Womelsdorf, Valiante et al. 2014, Urban-Ciecko and Barth 2016, Funk, Peelman et al. 2017).

#### 3.2.3 Cortical column communication through VIP-controlled disinhibition

The disinhibition of cortical PCs gates information flow through and between cortical columns (Walker, Möck et al. 2016). One of the factors underlying this PC disinhibition is the inhibition of SST cells during active cortical processing, which enhances distal dendritic excitability (Gentet, Kremer et al. 2012). The activation of VIP neurons strongly inhibits dendritic-targeting SST interneurons mediating surround inhibition, which leads to PCs disinhibition (suppress the inhibition on PCs) (Lee, Kruglikov et al. 2013, Pi, Hangya et al. 2013, Fu, Tucciarone et al. 2014, Pfeffer 2014, Yang, Murray et al. 2016). This disynaptic mechanism of disinhibition seems to be a generic motif able to suppress the blanket of inhibition mediated by SST neurons (Fino and Yuste 2011, Karnani, Agetsuma et al. 2014). It has been demonstrated indeed in motor, sensory and associative neocortical areas that transient SST activity suppression by VIP activation occurs during visual processing (Lee and Mihalas 2017), somatosensory integration (Lee, Kruglikov et al. 2013, Sohn, Okamoto et al. 2016), locomotion (Dipoppa, Ranson et al. 2018), top-down modulation (Ayzenshtat, Karnani et al. 2016) and plasticity during perceptual learning (Williams and Holtmaat 2018). Since VIP neurons are targeted by long-range cortical glutamatergic projections, they represent a key factor for distal cortico-cortical activation through disynaptic disinhibition.

#### 3.2.4 Formal description of neocortical model

Based on the above information, we have developed a neocortical module involving PCs and three types of inhibitory subpopulations, namely, BC, SST and VIP (see **Error! Reference source not found.**B, left panel). BC and SST receive excitatory inputs from PCs that are reciprocally inhibited by both of them. Pyramidal collateral excitation was also implemented *via* an excitatory feedback loop passed by a supplementary excitatory population (PC’) analogous to PC, except that it projects only from and to subpopulation PC. The electrical gap-junction mentioned in section 3.2.1 was implemented through an inhibitory feedback loop characterized by a connectivity constant 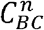, where *n* is the index of the NMM. Communication through disinhibition mentioned in section 3.2.3 was modeled by inhibitory projections; first from VIP to SST, and second, from the latter to BC. The nonspecific influence from neighboring and distant populations was modeled by a Gaussian input noise corresponding to an excitatory input 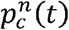 that globally describes the average density of afferent action potentials. The set of ordinary differential equations (ODEs) modeling the neocortical module is included as a Supplementary Material (see Supplementary section 1.1).

### 3.3 A local neural mass model of thalamic activity

The thalamus is considered as a complex relay extensively connected with the cortex, as well as most subcortical areas. This central position underlies its key role in several cognitive functions including perception, attention, memory and consciousness. Importantly, even a limited damage in the thalamus can have major consequences on all the aforementioned functions (Ward 2013). The thalamo-cortical circuitry as a neural correlate of consciousness has been mentioned by consciousness theories. More evidence for this role was provided by a study analyzing the metabolic activity of posterior midline cortical areas driven by the thalamic nuclei, across different altered consciousness states (Laureys, Boly et al. 2006). The authors reported that unresponsive wakefulness syndrome (UWS) patients can be differentiated from minimally conscious state (MCS) patients by a difference in glucose metabolism in these areas. The pivotal role of thalamo-cortical loop in the generation of slow waves and the up-and-down state was appraised in the review of Crunelli et al. (Crunelli, David et al. 2015) where they enumerated the main lines of evidence supporting this assertion, namely, the strong interconnection between thalamic cells (TCs) and neocortical layers involved in slow waves suggesting that thalamic nuclei can control up-and-down state dynamics in neocortical circuits, the early TC firing in relation to the initiation of cortical UP states, the rhythmic up-and-down state generated by TC neurons and TRN in isolated conditions, and the neocortical UP states readily induced in head-restrained mice by selective optogenetic activation of TC neurons proving the stimuli role of the latter.

The importance of thalamo-cortical connectivity has motivated the development of thalamo-cortical models simulating the interactions between the cortex and thalamus at a mesoscopic level (Suffczynski, Kalitzin et al. 2004, Sotero, Trujillo-Barreto et al. 2007, Bhattacharya, Coyle et al. 2011, Roberts and Robinson 2012, Mina, Benquet et al. 2013, Sen Bhattacharya, Cakir et al. 2013, Cona, Lacanna et al. 2014). While some models were developed to generate alpha activity (8-12 Hz) (Sotero, Trujillo-Barreto et al. 2007, Bhattacharya, Coyle et al. 2011, Sen Bhattacharya, Cakir et al. 2013), others were used to simulate the sleep-wake cycle (Suffczynski, Kalitzin et al. 2004, Roberts and Robinson 2012, Cona, Lacanna et al. 2014).

Since our aim is to develop a computational model able to reproduce brain rhythms corresponding to different consciousness states (e.g. sleep-wake cycle), the inclusion of the thalamus in the model is crucial. A description of the thalamus model is provided in **Error! Reference source not found.**.B. The thalamic module includes one population of excitatory glutamatergic neurons TCs, and two inhibitory interneurons from the TRN, TRN_1_ and TRN_2_ accounting for fast and slow GABAergic IPSPs, respectively. TCs receive GABAergic IPSPs with slow and fast kinetics from the TRNs, whereas the latter receive excitatory inputs from the former. Similarly to the cortical module, a Gaussian input noise corresponding to excitatory input 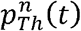 was used to represent nonspecific inputs on TCs. The set of ODEs modeling the thalamic module is provided as a Supplementary Material (see Supplementary section 1.2).

### 3.4 Modeling of large-scale cortico-cortical and thalamo-cortical connectivity

#### 3.4.1 Cortico-cortical connections

PCs originating from a single cortical column target several cell types in distant cortical columns. Glutamatergic PCs target not only remote PCs by common feedforward excitation, but also GABAergic cells by disynaptic cortico-cortical feedforward inhibition (FFI)(see Figure 2). Even if PV^+^ BCs have been shown to receive stronger thalamocortical and intracortical excitatory inputs than SST neurons, it appears that cortico-cortical FFI could be mediated by both types of interneurons (Ma, Liu et al. 2010, Tremblay, Lee et al. 2016). Importantly, cortico-cortical glutamatergic axons also target VIP neurons (Sohn, Okamoto et al. 2016).

**Figure 2.**
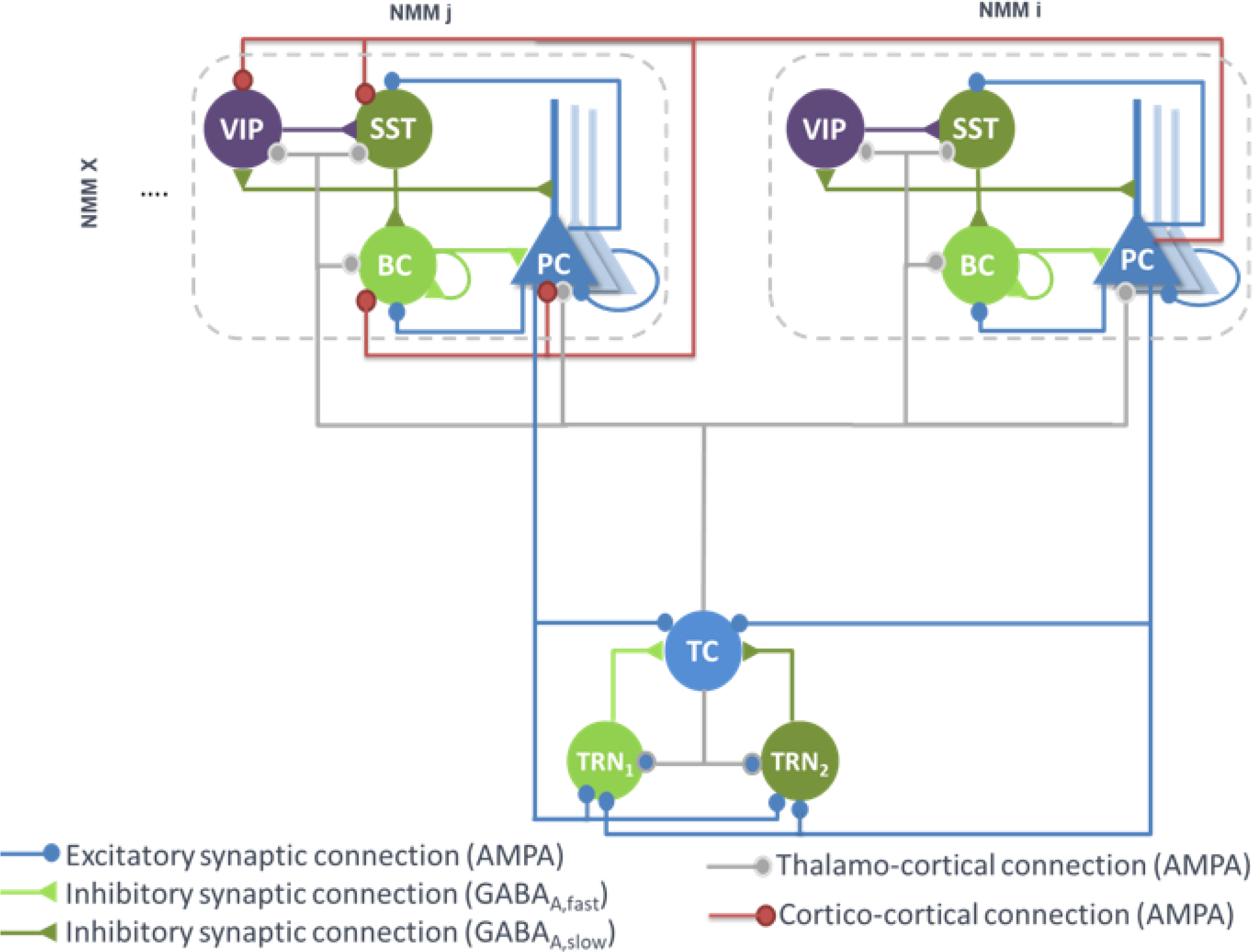
Large-scale architecture of the model. Illustration of the synaptic projections between cortical modules, and between thalamic and cortical modules. Note the presence of long-range thalamocortical and cortico-cortical feedforward inhibition. For the sake of clarity, long-range connections between cortical NMM_i_ and NMM_j_ are unidirectional, whereas in the model, pyramidal cells of NMMj also project on neurons of NMM_i_. The strengh of the synaptic input of PC onto distant VIP cells is larger than the input onto the other distant interneurons (SST, Basket).

Therefore, feedforward excitation was included in the model by means of a connectivity constant 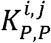 modeling the average strength of glutamatergic projections from pyramidal subpopulations in NMM “*i*” to their counterpart in NMM “*j*”. Disynaptic cortico-cortical feedforward inhibition was similarly integrated *via* the connectivity constants 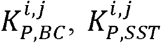 and 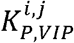 denoting glutamatergic projections from PC subpopulations in NMM “*i*” to BC, SST and VIP in NMM “*j*”, respectively. The set of ODEs modeling the cortico-cortical connections is included as a Supplementary Material (see Supplementary section 1.3, equations (9)-(12)).

In long-range cortico-cortical connections, the time-delay between NMMs “*i*” and “*j*” was controlled by a distance parameter *D*^*i, j*^ that reports the Cartesian distance in centimeters (cm) between the two populations. By setting the conduction velocity of action potentials in the brain to *c* (*c* ∈ [10, 100] cm/s), the time-delay taken by NMM “*j*” to receive a firing rate is straightforwardly deduced as *D*^*i, j*^/ *c*.

#### 3.4.2 Thalamo-cortical connections

The main connections between the thalamus and neocortex were taken into consideration in the model (see Figure 2). As in classical thalamocortical models, TCs receive glutamatergic excitatory postsynaptic potentials (EPSPs) from PCs. In turn, PCs receive excitatory input from TCs. Similarly, TRNs also receive excitatory cortical projections. In terms of GABAergic cortical targets, thalamic projections mainly target PV^+^ basket cells (Cruikshank, Lewis et al. 2007, Yang, Carrasquillo et al. 2013). In the adult brain, thalamic projections onto SST neurons are present but are much weaker as compared to the projections onto PCs and PV^+^ cells (Ji, Zingg et al. 2016). However, robust thalamocortical activation of non-Martinotti dendritic-targeting GABAergic interneurons has been demonstrated (Tan, Hu et al. 2008). Long-range connections from cortical areas and/or thalamic nucleus areas can activate VIP neurons, which in turn inhibit SST neurons, and disinhibit PCs dendrites. Such dendritic disinhibitory circuit has been proposed to gate excitatory inputs targeting pyramidal dendrites (Yang, Murray et al. 2016, Williams and Holtmaat 2018). Note that the thalamic compartment implements FFI, since it induces first an EPSP (PCs activation) followed later on by an IPSP (cortical interneurons activation).

Since only one thalamic population was considered in this model, the TC projections to - and from - PCs were included in the model. Connectivity constants 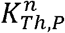 and 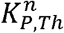, allow to adjust the strength of efferent TC projections to PCs of NMM “*n*” and efferent pyramidal projections from the NMM “*n*” to TC, respectively. Projections from PCs of NMM “*n*” onto TRN_1_ and TRN_2_ were also integrated and adjusted with the connectivity constants 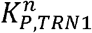 and 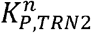, respectively. Likewise, projections from TC to GABAergic interneurons in NMM “*n*”, namely BCs, SSTs and VIPs, were included through the connectivity constants 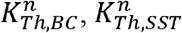 and 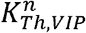 respectively. The set of ODEs modeling the thalamo-cortical connections is described in Supplementary section 1.3 (equations (13)-(19)). It is noteworthy that time-delays between the thalamus and the cortical NMMs were included as in the cortico-cortical long-range connections.

### 3.5 Implementation and parameter tuning

An important step in NMM approaches consists in tuning the model parameters. Some of these parameters, namely time constants in the three modules, were set close to the “standard values” used in neuronal population models, while other parameters such as connectivity constants were adjusted according to the target EEG activities. Two classical well-known examples of conscious and unconscious states are deep sleep- a.k.a. slow wave sleep (SWS) characterized by delta oscillations (0-4 Hz), and wakefulness (background activity). The objective was to reproduce these manifestations of consciousness modulation in the model.

In Supplementary Table 1, we provide physiological interpretation and values of model parameters. Regarding the model output, the signal simulated at the level of PCs in the cortical compartment was chosen as the model output since it corresponds to the sum of PSPs, which is the main contribution to LFPs recorded in the neocortex.

When several cortical NMMs are interconnected, a simple way to handle large-scale connectivity is to arrange the connectivity constants in arrays where line and column indices refer to source and target NMMs, respectively. Based on section 3.4, there are two categories of connectivity array, “excitatory to excitatory” and “excitatory to inhibitory” arrays. In the first category, we consider the matrix ***K***_***EXC***_ whose (*i*, *j*)^th^ element represents the glutamatergic projections from NMM “*i*” onto an excitatory subpopulation in NMM “*j*”. Hence, if both of them belongs to the cortical module, the (*i*, *j*)^th^ element would correspond to 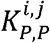. However, if NMM “*i*” coincides with a thalamic population, the (*i*, *j*)^th^ element would be then equal to 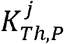. In the second category, a similar scheme is considered with the difference that the target subpopulation is always an inhibitory interneuron (BC, SST, VIP or TRNs). Consequently, fives arrays are considered, namely, ***K***_***BC***_, ***K***_***SST***_, ***K***_***VIP***_, 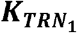 and 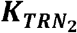 (see Supplementary Figure 1). Note that when only one thalamic NMM is considered, the two last matrices are reduced to vector arrays. Cortico- and thalamo-cortical time-delays were similarly arranged in matrix ***D*** whose (*i*, *j*)^th^ element coincides with the aforementioned *D*^*i, j*^.

The set of second order stochastic nonlinear ODEs obtained for all synaptic interactions present in the model was numerically solved using a fixed step (Δt = 1 ms) 4^th^-order Runge-Kutta method. The model was implemented using an object-oriented language (Objective-C).

To study the model behaviour, we followed a two-step approach. We first implemented a “toy model” with small number of coupled neural masses. This reduced-complexity model allowed to assess the effects of cortico- and thalamo-cortical connectivity matrices on cortical activities associated with different consciousness states (see sections 3.2, 3.3 and 3.4). After validation in the toy model, all mechanisms were implemented in an extended, more realistic model, where the whole brain was considered.

#### 3.5.1 Toy model of cortical activity

The toy model was composed of one thalamic population connected to four cortical populations (*N* = 4) that were identically tuned. The default values of NMMs intrinsic parameters are listed in Supplementary Table 1. Unless explicitly mentioned, these parameters were kept unchanged for all neural activities generated afterwards (SWS and background activity). The objective was to verify, in a simplified model, the hypothesis that cortical activity is modulated from deep sleep to wakefulness by tuning only thalamo- and cortico-cortical connectivity, so that, when thalamocortical connectivity increases, the model goes deeper into sleep. This connectivity mechanism was implemented in all connectivity matrices in the model (see Supplementary Figure 2).

#### 3.5.2 Whole brain model: from cortical activity to EEG

The pipeline used for the simulation of scalp EEG data is described in Figure 3. In order to obtain a “realistic” activity during wakefulness and SWS over the entire neocortex, we considered one thalamic population and 66 cortical populations (*N*+1 = 67). Each time-course at the output of these populations represented the activity of one macro-region of the anatomical parcellation described in (Desikan, Segonne et al. 2006), in which the activity was assumed to be homogenous. In order to generate 67 time-courses from the coupled NMMs (66 cortical regions plus the thalamus), we used a combination of connectivity arrays (Figure 3B). First, a matrix of connection weights ***K***_***DTI***_, representing a density of fibers between all pairs of 66 cortical regions was used to set the structural connectivity. This matrix, obtained from DTI, is provided in (Hagmann, Cammoun et al. 2008). Second, we considered additional functional horizontal (i.e. cortico-cortical) matrices ***K***_***Hx***_, reproducing the coefficients weights used for both wakefulness and sleep in the toy model, taking also into account the new number of populations. In order to apply the connectivity weights defined earlier in the toy model only to pairs of NMMs that are structurally connected. Structural and horizontal functional matrices were combined using the Hadamard product. Finally the vertical (i.e. thalamo-cortical) connectivity ***K***_***Vx***_, was added to this product to obtain a set of anatomo-functional connectivity matrices 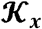 such that

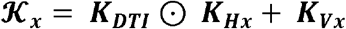

with *x* = {*EXC*, *SST*, *BC*, *VIP*, *TRN*1, *TRN*2}. All connectivity matrices are described in Supplementary Figure 3.

**Figure 3.**
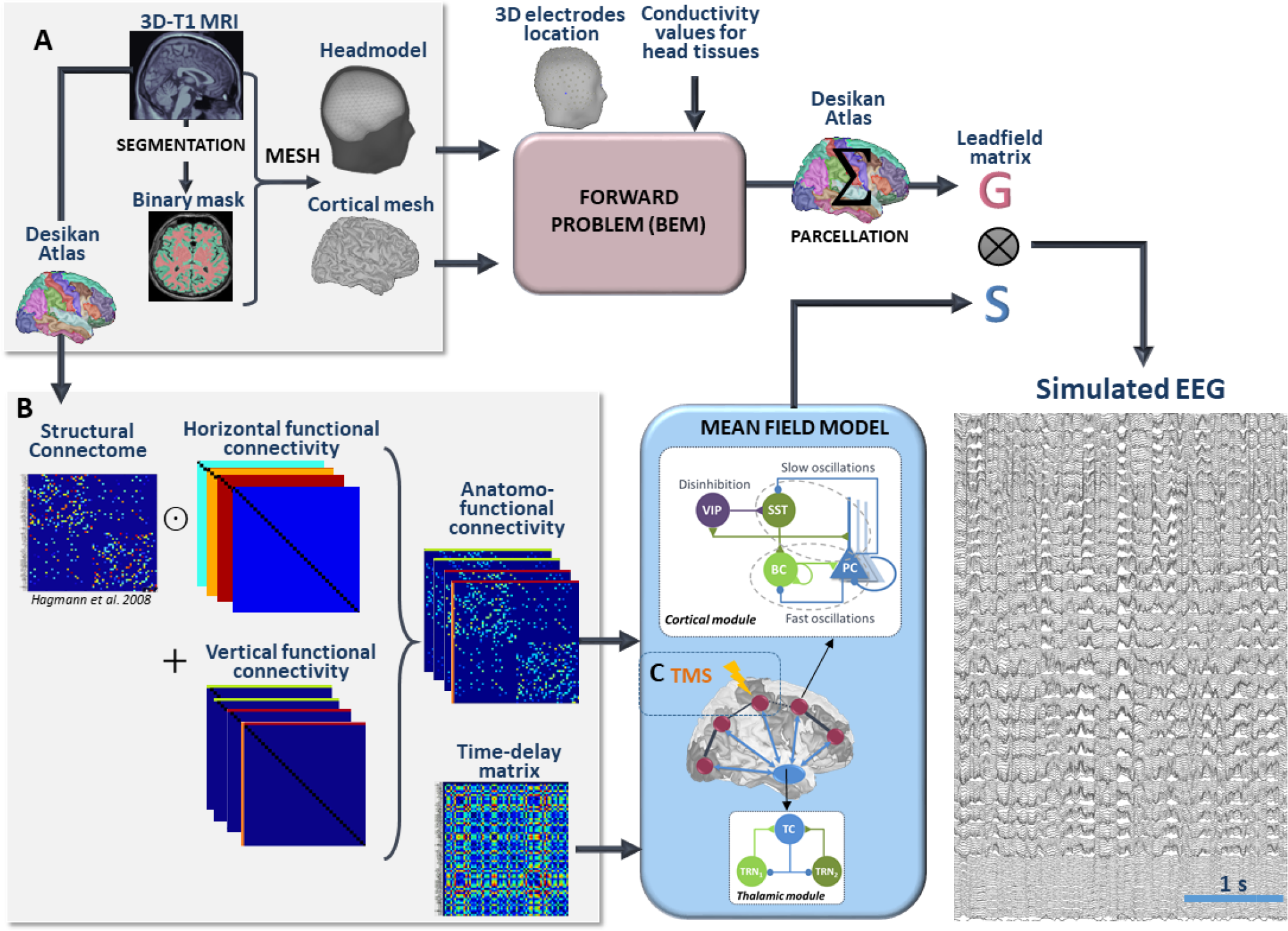
Full processing pipeline leading to the simulation of scalp EEGs. The pipeline to simulate EEG is a two-step process. **(A)** First, the forward problem is solved at the level of 257 scalp electrodes from a dipole layer constrained to the surface of a cortical mesh (15000 vertices) midway between the Grey/White matter interface and the cortex surface. The boundary Element Metho is used for the calculation within a realistic head model that accounts for the conductivity properties and the geometry of brain, skull and scalp. This step provides a 257×150000 leadfield matrix ***A*** representing the contribution of each individual cortical dipole at each of the 257 scalp electrodes. In this matrix, leadfield vectors belonging to a common region of the Desikan Atlas are added to obtain a simplified 66 × 257 matrix **G**. Second, the time courses ***S*** at the whole brain level are obtained in the mean field model from a set of 66 cortical and one thalamic coupled NMMs. **(B)** Coupling between these 67 NMMs is done using combination of connection weight matrices. Pairs of structurally connected cortical NMMs are first defined from a matrix of connection weights representing a density of fibers between all pairs of 66 cortical regions of the Desikan Atlas. This matrix is provided in (Hagmann, Cammoun et al. 2008). Using an element-wise multiplication, this matrix is combined with a set of horizontal (i.e. cortico-cortical) functional connectivity matrices that reproduce the coefficients weights used for wakefulness and sleep in the toy model. Vertical (i.e. thalamo-cortical) connectivity matrices are added to each of these products to obtain connectivity weight matrices that account for anatomical connections as well as cortico-cortical and thalamo-cortical connectivity matrices. Cortico- and thalamo-cortical time-delays were similarly organized in the form of matrices where the elements represent the Cartesian distance between cortical NMMs divided by the mean velocity of travelling for action potentials. **(C)** The mean field model includes explicitly the contribution of an external stimulus term that represents the effect of TMS. At the output of the pipeline, scalp EEG signals at the level of 257 channels are obtained as the produc leadfield ***G*** and source time courses ***S***.

Using this combination of connectivity weights, and specific parameters defined in Supplementary Material for delta and background activity, we built a spatio-temporal source matrix ***S*** containing the time-varying activities of the thalamus and of all cortical macro-regions.

To reconstruct simulated scalp EEG data, we first solved the forward problem using the Boundary Element Method (BEM, OpenMEEG, (Gramfort, Papadopoulo et al. 2010). To this end, a realistic head model was built in Brainstorm (Tadel, Baillet et al. 2011) from the segmentation of a template T1 MRI (Colin 27 template brain, (Holmes, Hoge et al. 1998)) previously obtained using the Freesurfer image analysis suite (http://surfer.nmr.mgh.harvard.edu/, (Dale, Fischl et al. 1999, Fischl, Sereno et al. 1999)), as illustrated in Figure 3A. The head model consisted in three nested homogeneous mesh surfaces shaping the cortical surface (642 vertices), the skull (642 vertices) and the scalp (1082 vertices) with conductivity values of 0.33 Sm^−1^, 0.0082 Sm^−1^ and 0.33 Sm^−1^, respectively (Goncalves, de Munck et al. 2003). The forward problem was then numerically calculated for each vertex of the source mesh obtained from the segmented white matter/ grey matter interface of the same template brain. As a result, the leadfield matrix ***A***, represented the contribution of each dipole of the source mesh at the level of 257 scalp electrode positions (high density EEG), placed over the scalp according to the geodesic convention (EGI®, Eugene, USA). All leadfield vectors of ***A*** belonging to a common region of Desikan Atlas were added to obtain a simplified 66 × 257 matrix ***G***. The spatio-temporal matrix ***X*** of simulated EEG data was given by the matrix product:

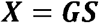

The entire pipeline enabling the simulation of scalp EEG is provided in Figure 3. Finally, as detailed in the Supplementary Material, a total number of 1060 ODE’s must be solved to run the model. To give an idea of the required computing time, 60 seconds of simulated EEG could be simulated in 49 seconds on a 3.5 GHz 6-Core Intel Xeon EG with 64GB of 1866 MHz RAM (OsX Mojave) and in 72 seconds on a standard Intel 2.5 GHz 2-Core Xeon EG with 8GB RAM (Windows 10). Therefore, the model performed almost real-time simulation on a standard PC.

### 3.6 Cortical response to TMS and PCI

In order to simulate TMS-evoked EEG responses that can be compared to those recorded experimentally (Casali, Gosseries et al. 2013), we included the effect of an exogenous, TMS-induced, stimulation in the whole-brain model. The model can simulate not only the activity of the 67 pre-defined anatomical regions, but also the EEG signals recorded by 257 scalp electrodes. Furthermore, since the model includes, as a connectivity matrix between the 67 regions, a DTI-derived connectivity matrix (Hagmann, Cammoun et al. 2008), it is possible to track the spatio-temporal dynamics of the stimulation-evoked network, i.e. the activated regions, along with the peak latency for each region. In addition, since the model can simulate “wake” and “sleep” states, it provides the opportunity to compare TMS-evoked responses in these two states of consciousness, which have been experimentally recorded in humans. Therefore, the structure of our computational model provides a unique framework to interpret TMS-evoked EEG responses obtained in humans.

In principle, TMS involves a high-intensity current flowing through the stimulation coil, thereby generating a magnetic field penetrating without attenuation through the head. Since the stimulation pulse is very short (≈0.1 ms), the magnetic field gradient dB/dt is extremely high (> 30,000 T/s), resulting per Maxw Ampere’s law into an electric field at the level of brain tissue. Since the electric field induced in brain tissue is high (> 100 V/m, (Miranda, Hallett et al. 2003)), this induces neuronal firing, presumably at the bending point of cortical axons, triggering a series of complex activations within the stimulated area (Di Lazzaro and Ziemann 2013).

We used a simple approach to represent the effect of TMS, consisting in simulating an afferent volley of action potentials (in terms of pulses/s) to the stimulated cortical region. Since the 1 ms time step used to numerically solve the equations of the model was higher than the duration of an actual TMS pulse, the simulated length of this volley of incoming action potential was adapted and fixed to 5 ms. The amplitude of the simulated evoked volley of action potentials was fixed to 1000 pulses/s, and was applied to each cellular subtype of the stimulated region (i.e., PCs and all types of GABAergic interneurons). For the purpose of this paper, we chose to simulate the stimulation of the right cuneus in both conditions (“wake” and “sleep”), which is known to have a number of anatomical connections that should result in the propagation of the TMS-evoked responses in several cortical structures. The repetition rate of the TMS protocol was fixed to 2 seconds, and the total simulation duration was a full minute, resulting in a total of 30 TMS-evoked responses available at the source level. The EEG activity at the level of each scalp electrode was then computed from the simulated source activity using our EEG forward problem pipeline, as described previously. From the simulated TMS-evoked responses, the studied outcomes were the anatomical regions activated by TMS, and also their latency from the onset of the TMS pulse.

Finally, simulated EEG responses were used to compute the Perturbational Complexity Index (PCI) (Casali, Gosseries et al. 2013). PCI is a measure of TMS-evoked responses complexity, based on the Lampel-Ziv compression algorithm. The basic idea is that, if a sequence of information is complex, then it can be only marginally compressed (low compression rate); and on the opposite a very simple sequence can be described by a very limited amount of information (high compression rate). To derive PCI, a process similar to the one proposed by Casali et al. was used (Casali, Gosseries et al. 2013):

- The sources activity is stored in a 2D matrix (number of lines equal to the number of channels, number of columns equal to the number of time points).
- A threshold value for the simulated sources activity was set by preserving the highest 20% (proportional threshold) of values once fixing to 0 values below it.
- Each value of the sources activity matrix that is above or equal to the threshold value is set to 1, while all other values are set to 0 (binarization process).
- The Lempel-Ziv algorithm is then applied to the resulting binary matrix.

PCI values were computed in the two different scenarios, corresponding to the “wake” and “sleep” state, respectively.

Overall, our implementation of TMS-evoked responses enables a meaningful comparison with human data, since it results in similar experimentally measurable quantities: impacted anatomical regions, latency of the TMS-evoked response within specific regions, and complexity of the brain-scale response through PCI.

## 4 Results

### 4.1 Toy model: the impact of cortico- and thalamo-cortical connectivity on the cortical rhythms during sleep and wakefulness

During the deep sleep state, the thalamocortical connectivity is meant to be strong compared to the cortico-cortical one (see Supplementary Figure 2, first column). Our strategy consisted in progressively and simultaneously decreasing and increasing the thalamo- and cortico-cortical connectivity, respectively, in order to switch to wakefulness (see Supplementary Figure 2, column 2). Note that this connectivity process was mostly reflected by arrays and ***K***_***EXC***_ and ***K***_***BC***_, to simulate the strong thalamic projections onto PCs and BCs as reported in section 0. It is also noteworthy that no time delays were injected in the toy model example.

In Figure 4, we provide an example of simulated LFPs a.k.a. intracerebral EEG (iEEG) in comparison with real (human) ones. The left column depicts the scheme of high (and low) thalamo- (and cortico-) cortical connectivity. As depicted, the reinforcement of the thalamocortical loop (TC→PC→TRNs and TC→BC) resulted in the generation of delta oscillations that characterize SWS. In this example, the simulated delta was around 2-3 Hz (slightly faster than real iEEG). Conversely, by reducing the thalamocortical loop (right column), delta waves disappeared and were replaced by background activity, indicative of oscillatory changes observed during the switch from sleep to wake. We emphasize that only the large scale connectivity was tuned while all remaining parameters were kept unchanged, confirming the crucial role of thalamo- and cortico-cortical connectivity in modulating consciousness.

**Figure 4.**
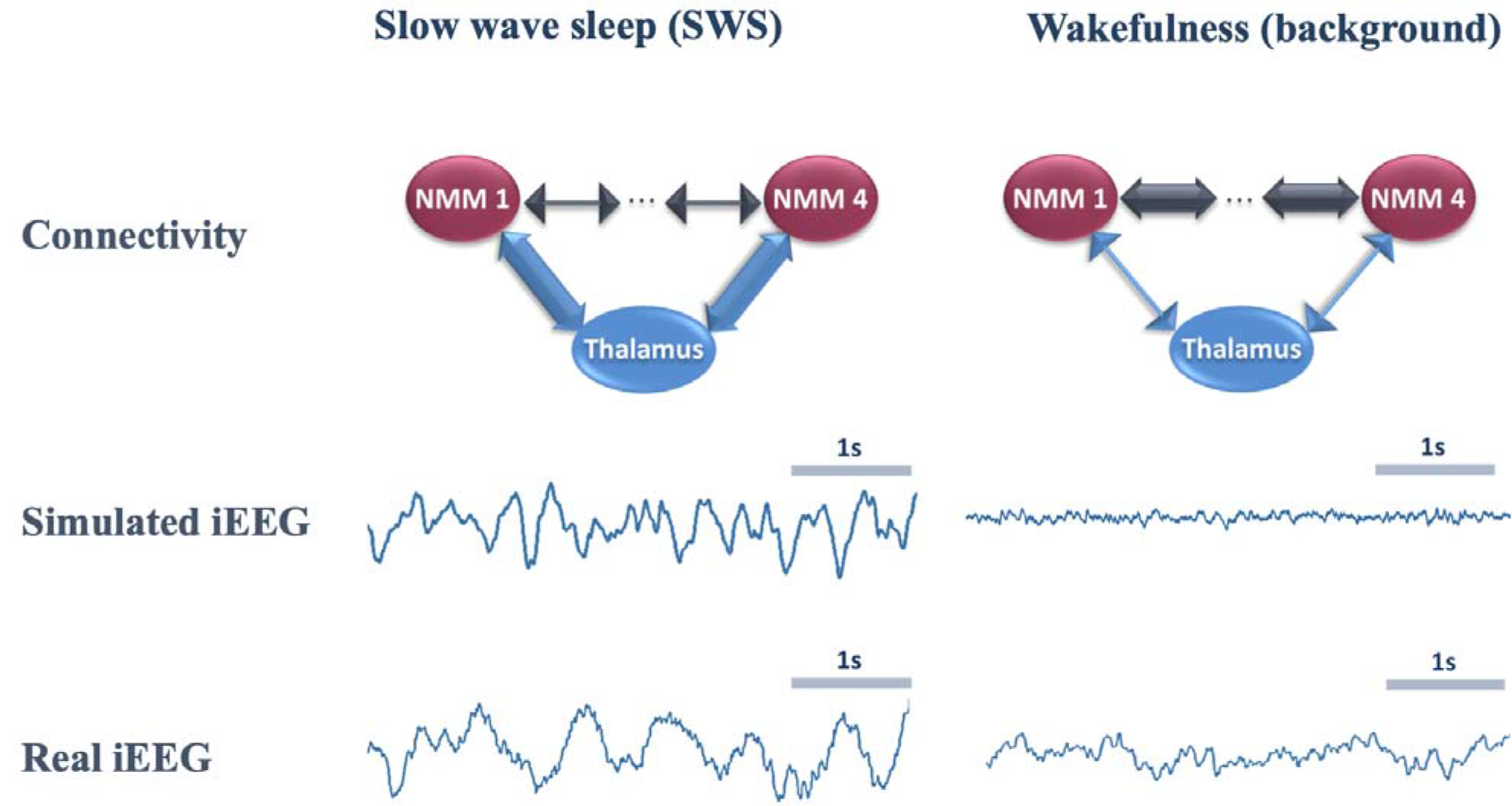
Comparison of real and simulated intracerebral EEG (iEEG) (toy model, *N* = 4). *Left column*: In the condition of high thalamo-cortical connectivity (i.e. low cortico-cortical connectivity), signals generated by the mean field model are characterized by delta waves (~ 4 Hz). These simulated signals are similar (although slightly faster) to delta waves recorded by intracerebral EEG (iEEG) during SWS in non-epileptic cortical regions of one patient undergoing an invasive EEG exploration. *Right column*: In the condition of low thalamo-cortical connectivity, signals generated by the mean field model are similar to background activity recorded by iEEG in real conditions during wakefulness. Note that these iEEG recordings are performed in patients who are candidate to epilepsy surgery. For the sake of this study, only iEEG signals that do not show epileptic activity were retained.

### 4.2 Whole brain model: the impact of cortico- and thalamo-cortical connectivity on scalp EEG rhythms during sleep and wakefulness

The morphology of simulated intracerebral signals was not modified when connectivity matrices were adapted to a larger number of NMMs in order to account for whole-brain activity. In the case of low thalamo-cortical connectivity (wakefulness condition), background activity was similar to signals obtained in the toy model and resembled real intracerebral background activity recorded during wakefulness in humans. Similarly, by increasing thalamo-cortical connectivity, the whole-brain model generated delta activity at a mean frequency of 3.8 Hz that was consistent with the morphology and spectral content of delta activity obtained both from the toy model and from real intracerebral recordings during SWS.

Signals obtained with the whole-brain model in both conditions of wakefulness and SWS were used in the forward calculation to generate simulated scalp EEG data at the level of 257 electrodes (Figure 5). In the low-thalamocortical connectivity condition, simulated scalp EEG (Figure 5A) resembled scalp EEG background activity as recorded in humans during wakefulness (Figure 5C). The spectral analysis disclosed similar sub-band distribution in the simulated *vs*. real case, although simulated signals contained more beta frequency than real background activity. In the high-thalamo-cortical connectivity condition, simulated scalp EEG (Figure 5B) was comparable to scalp EEG signals recorded in humans during SWS (Figure 5D). In the simulation case, the peak frequency was 3.8 Hz, thus slightly higher than in the real case (2 Hz). Topographical voltage maps at the peak of delta waves showed analogous distribution over the vertex, the activity in the simulated case being slightly more posterior than on the real case example.

**Figure 5.**
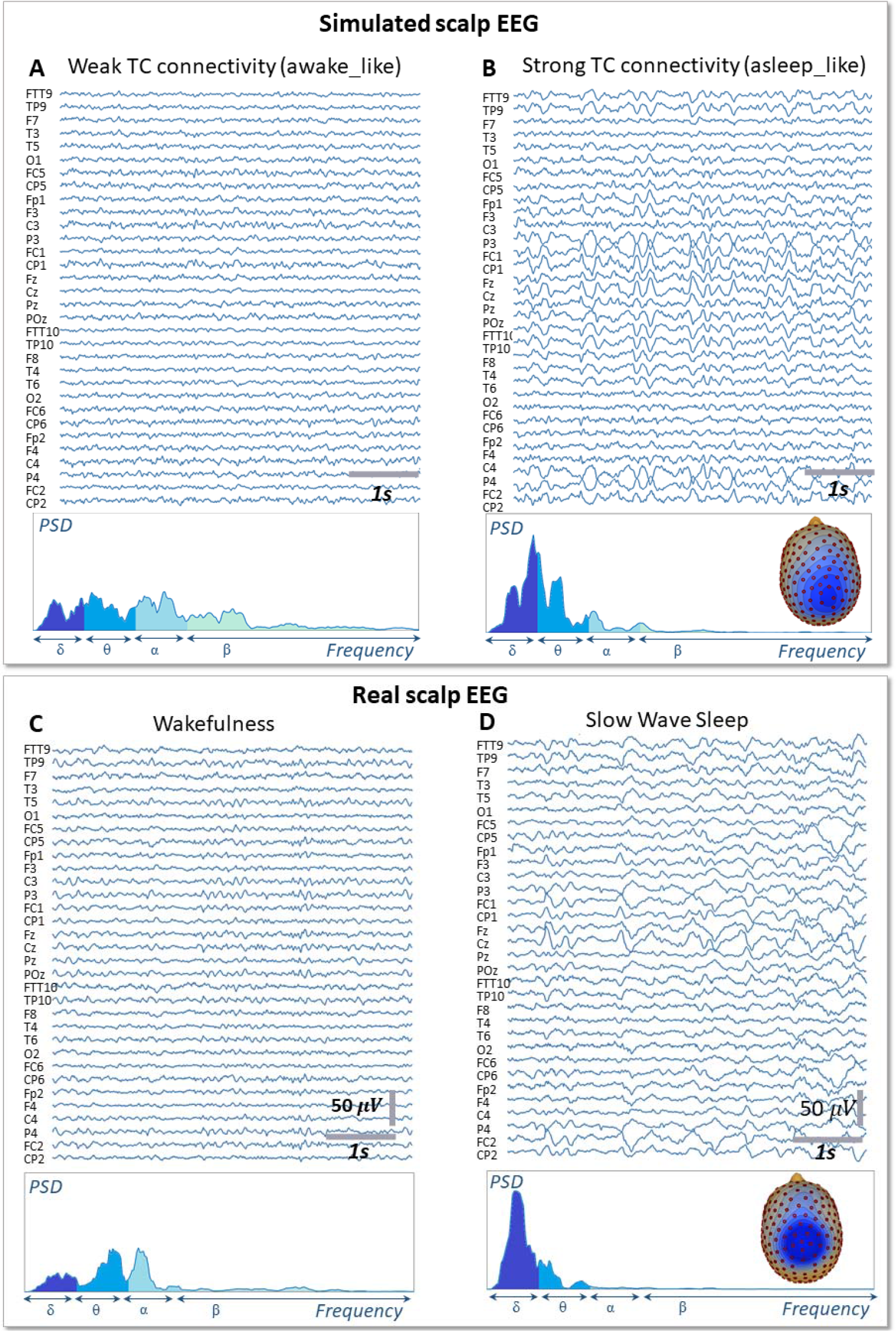
Simulated vs. Real EEG during wakefulness and SWS. Signals simulated with the whole-brain model using weak thalamo-cortical connectivity parameters **(A)** display background activity. The morphology and spectral content of these simulated signals is similar to scalp EEG recorded in a human subject during wakefulness in humans **(C)**, except for a higher power spectral density in the beta sub-band. Signals simulated with the whole-brain model using strong thalamo-cortical connectivity parameters **(B)** display delta waves similar to the activity recorded in real condition during SWS in humans **(D)**. The spectral content of signals as well as the topographical voltage distribution at the peak of delta waves were similar in the simulated and real conditions.

### 4.3 Bridging brain circuits, TMS-evoked EEG responses and complexity metrics

In Figure 6, we describe the process used to estimate the complexity of TMS-evoked EEG responses within our brain-scale model and in humans (Casali, Gosseries et al. 2013). As depicted, the simulated TMS-evoked EEG response (Figure 6B) was very similar to the human response (Figure 6A), not only in terms of length (250 and 350 ms, respectively) but also in term of regions distant from the right motor cortex (the stimulated area) that are activated post-stimulation. Indeed, in the simulated and experimental data, a first activation of the right pre-central gyrus occurred within 15 ms from the TMS pulse, followed by activity evoked notably in the right precuneus within 40 ms and a common propagation in the contralateral left precuneus at about 60 ms from the stimulus. In both simulated and experimental TMS-evoked responses, activity was evoked in the right frontal lobe within 110-120 ms, with activation of the right superior parietal cortex within 175 ms (human data) and 150 ms (model data). The most important difference is that the human TMS-evoked response was notably longer as compared to the model.

**Figure 6.**
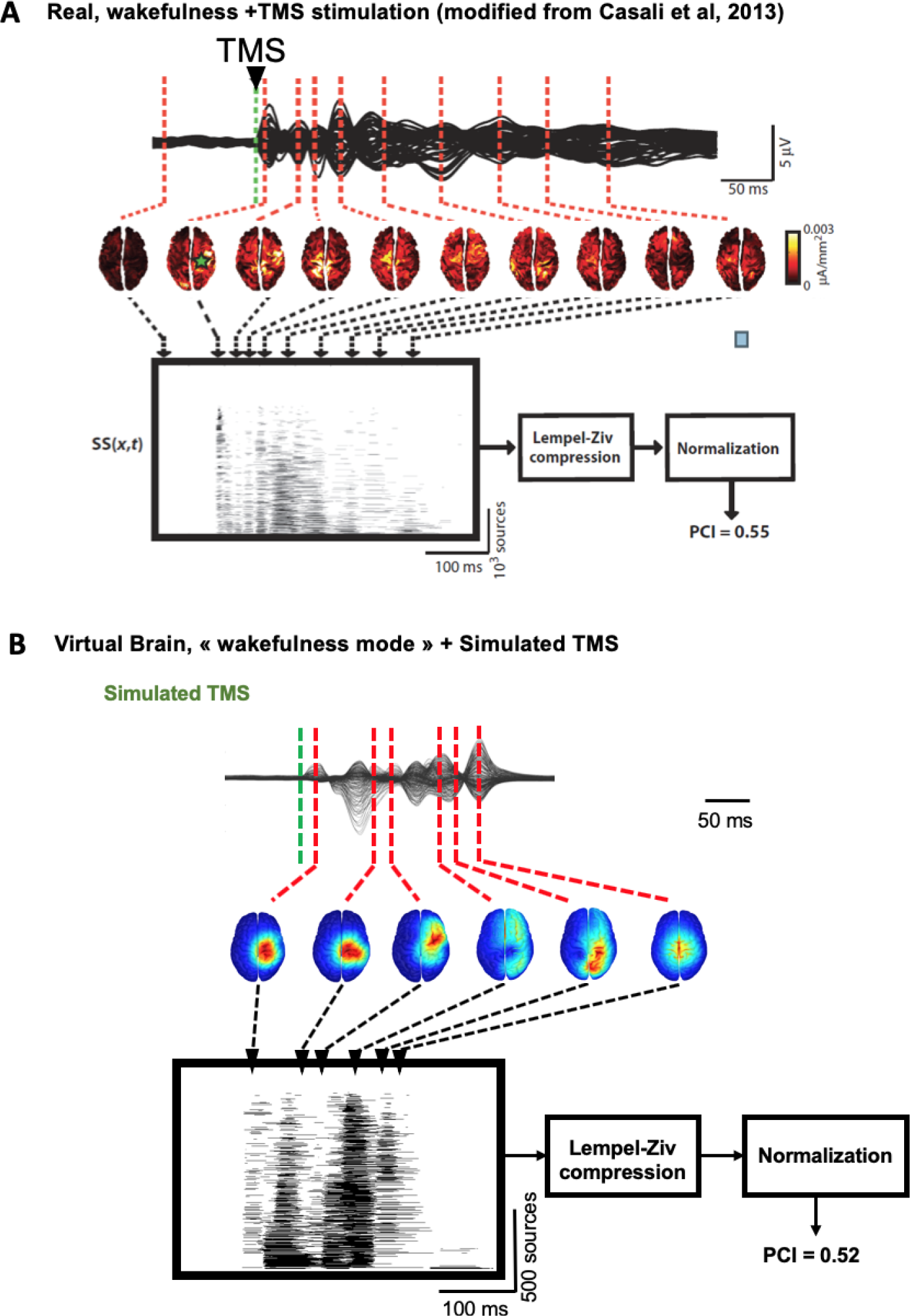
Comparison of TMS-EEG evoked responses *in silico* and in humans. **(A)** Time course of a human TMS-EEG response (modified from Casali et al., 2013) following stimulation of the motor cortex during wakefulness. Once that cortical sources have been computed from EEG recordings, a spatio-temporal matrix of significant sources was built and the Lempel-Ziv compression algorithm was used to compute the complexity of the evoked response (Perturbational Complexity Index, PCI). (**B)** Time course of a simulated TMS-EEG response using our brain-scale, following stimulation of the motor area in the wakefulness mode. Cortical sources were reconstructed from the simulated EEG, and a similar procedure was used to compute PCI. A similar PCI value was obtained in the simulated and experimental TMS-evoked EEG responses in the awake state.

We then compared the simulated and experimental TMS-evoked responses in wakefulness and in sleep, along with their PCI value, which is presented in Figure 7. In the two cases, the right motor area is stimulated. In the case of wakefulness, as also illustrated in Figure 6, there was a satisfactory agreement in the duration and global shape of the response, and also in terms of the sequence of activated brain regions. More precisely, by comparing Figure 7A (upper) and B (upper), we can conclude that the tracking of the propagation of the TMS-evoked activity revealed that this propagation occurred, as expected, along documented inter-regional connections from the DTI-derived connectivity matrix used in the model. PCI values obtained were also extremely similar (0.52 and 0.51 for the model and for humans, respectively). Conversely, in the “Sleep” condition, the time course of the TMS-evoked response was significantly shorter (less than 200 ms), which is also comparable to TMS-EEG human recordings. In addition, another striking similarity with human data is that the TMS-evoked activity remained confined to the stimulated area, i.e. the right motor area. The PCI value in this condition was 0.19, which is very similar to the value obtained in humans during sleep (0.23, see Figure 7A, lower panel), notably lower than the “Wakefulness” condition, which is also consistent with human data (Casali, Gosseries et al. 2013). Therefore, despite using the exact same TMS pulse characteristics within the two conditions (“Wakefulness” and “Sleep”), the simulated response was drastically different within the model: in wakefulness, the TMS-evoked response resulted in a complex sequence of successive activations within distant, anatomically connected areas in ipsilateral and contralateral regions; while during sleep the TMS-evoked activity remained confined to the stimulation site, even when the anatomical connections were present as in the “sleep” condition.

**Figure 7.**
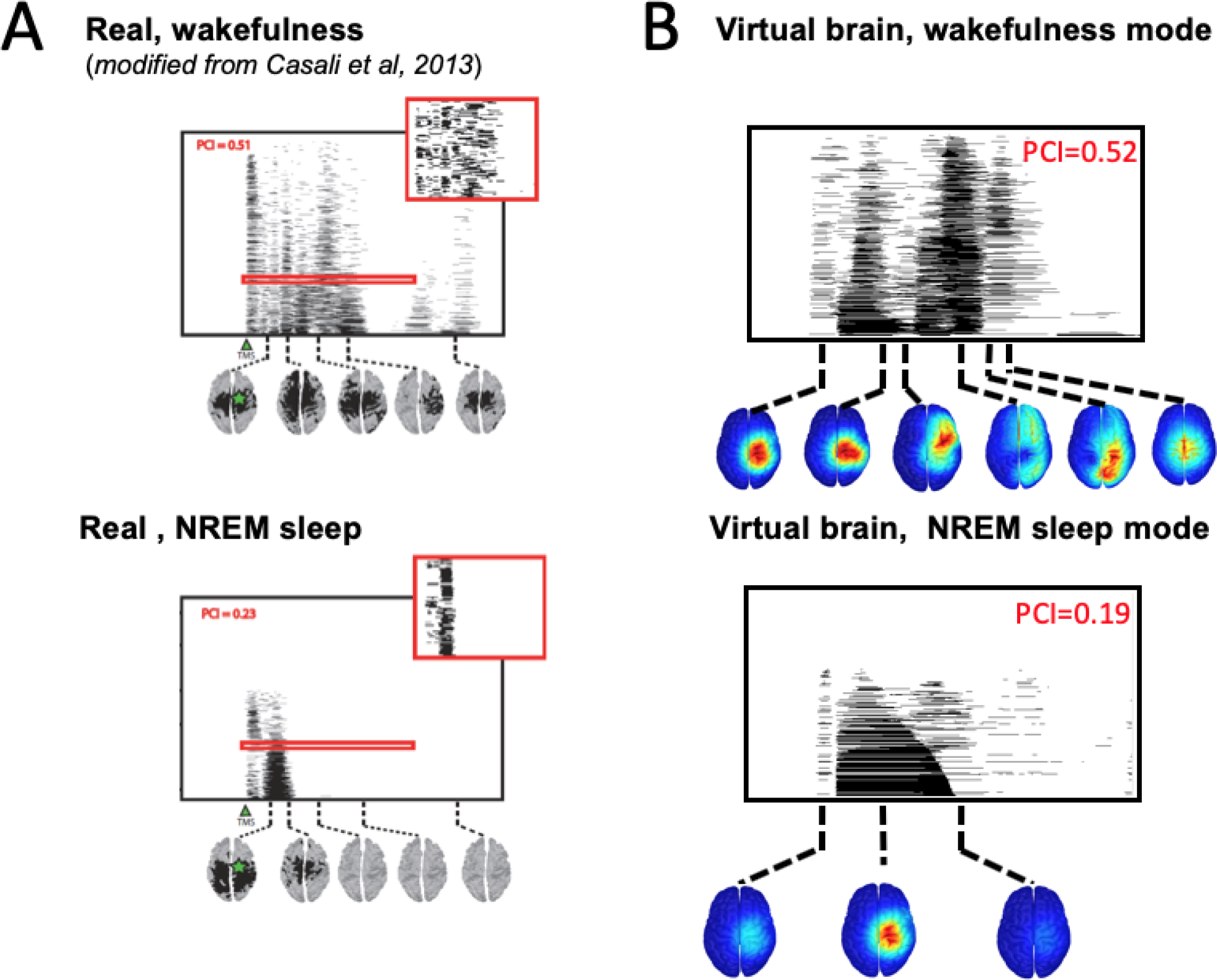
Comparison of TMS-evoked EEG responses in wakefulness and sleep. **(A)** TMS-evoked EEG responses obtained through TMS of the motor cortex in humans (upper panel, during wakefulness; lower panel, during sleep) with associated PCI values. (**B)** TMS-evoked EEG responses obtained through simulated TMS of the motor region (upper panel, in the wakefulness mode; lower panel, during the sleep mode) with associated PCI values.

## 5 Discussion and perspective

In this paper, we have developed the first brain-scale computational model that can reproduce neuronal activity patterns associated with various conscious states, while accounting for key microcircuits at the cellular type scale. A major asset of the model is its strong link with recent neurophysiological and neuroanatomical data: the main cellular types are included (PCs and different types of interneurons), along with their recently elucidated connections that underlie the selective disinhibition of distant neural populations (through VIP to SST projections), realistic synaptic kinetics, large-scale structural connectivity obtained in humans through DTI and propagation delays between regions based on their spatial distance. Furthermore, since the model features detailed micro-circuits, DTI-derived brain connectivity matrix and large-scale macro-circuits results also provided a bottom-up description of TMS-evoked responses in human.

The model offers novel, key insights into the maintenance of the neuronal activity associated with conscious states. Interestingly, this model including a variety of cellular subtypes with accurate synaptic kinetics provides realistic electrophysiological signals both at the level of cortical sources and of the EEG. First, it explains how thalamo-cortical (vertical) connectivity is critically involved in the gating of cortico-cortical (horizontal) information propagation. If thalamo-cortical activity is indeed rhythmically patterned, the communication between cortical areas is disrupted due to the resulting rhythmic inhibition. This result is therefore in line with the “connectivity breakdown” observed during sleep (Esser, Hill et al. 2009, Ferrarelli, Massimini et al. 2010, Casali, Gosseries et al. 2013).

Crucially, the various conscious states simulated with the model can be tuned by modulating only the thalamo-cortical input, which supports our hypothesis that this represents the crucial control parameter for consciousness, i.e. the dimension of wakefulness. It can be seen essentially as the possibility for information processing to take place between spatially distant cortical areas. Since the propagation of activity is impaired at the cortical level during sleep (absence of wakefulness), no cortical processing can take place, which explains the absence of consciousness. Second, regarding cortico-cortical activity (horizontal connectivity), since the model reproduces with a satisfactory qualitative and quantitative agreement the TMS-evoked EEG responses observed in awake humans (Casali, Gosseries et al. 2013), this suggests that 1) our modeling hypotheses and choices appear sufficient to capture the essence of TMS-EEG responses; 2) TMS-EEG responses are mainly driven by the underlying structural connectome, which is in line with recent research pointing at the tight links between structural and functional connectivity (Avena-Koenigsberger, Misic et al. 2017); 3) this modeling approach could be used to assist in the interpretation of TMS-evoked EEG responses in DOC patients, since our model links explicitly the underlying connectivity with the observed TMS-evoked response. Finally, the model validates that, at the brain scale, the disynaptic disinhibition of distant pyramidal cells through the activation of VIP neurons is indeed an effective mechanism enabling the transmission of activity between a few cortical regions. The model therefore confirms the role of a cellular-scale micro-circuit that regulates brain-scale propagation of activity within the cortex.

Among the possibilities to improve the realism and predictive power of our brain-scale model, the most immediate would be the use of structural connectivity matrices averaged among a large number of healthy participants, such as those from the Human Connectome Project (see the link http://www.humanconnectomeproject.org). Furthermore, no regional specificities were accounted for between the 66 cortical regions included within the model, since we used standard parameter values for the synaptic gains (e.g., A, B and G) and the same within-population connectivity parameters. By doing so, we have assumed that the large-scale anatomic structure of brain connectivity and the cellular-scale micro-circuits included are the main factors explaining the resulting simulated EEG signals.

It should be mentioned that in the case of deep sleep simulated signals, the median frequency of delta activity was higher in the model (2-3 Hz) as compared to human data (~1 Hz). This discrepancy is explained by the fact that model does not implement the mechanisms underlying slow oscillations (~1 Hz) generated in cortical and thalamic networks, among which: i) the sequence of depolarizing periods followed by silent periods (up to 2 s) during up-and-down states as well as ii) GABA_B_-mediated pre-synaptic slow inhibition that also appears to play some role (see review in (Neske 2015)). Other limitations likely explain the moderate discrepancies between the simulated and experimental TMS-EEG responses in terms of latencies and localizations, such as the lack of asymmetry in connectivity weights (i.e., all connections were assumed bidirectional and identical).

The future prospects regarding our brain-scale EEG model are numerous: in terms of consciousness studies, the model could be used to study the mechanisms underlying the so-called “slow-wave activity saturation”, delta-band activity that appears when the blood concentration of anesthetics is increased (Ni Mhuircheartaigh, Warnaby et al. 2013), and that constitutes a solid marker of the conscious state. Furthermore, this study improves our understanding of active probing paradigms of brain circuits in DOCs, such as the PCI and paves the way toward the design of optimized stimulation-based metrics to measure consciousness. The model indeed allows to test *in silico* novel neuromodulation protocols based on TMS, transcranial direct current stimulation (tDCS) and transcranial alternating current stimulation (tACS), aiming at quantifying the level of residual consciousness in DOC patients. Beyond applications for consciousness, the model could be exploited to understand the detailed dynamics of TMS-EEG responses and their underlying mechanisms, or to shed light on the mechanisms underlying the generation and propagation of epileptiform activity.

## 7 Author Contributions

SB: model implementation, simulations, data analysis, manuscript write-up. JM: model design, simulations, data analysis, manuscript write-up. IM: simulations, data analysis, manuscript write-up. FW: model design, data analysis, manuscript write-up. PB: model design, data analysis, manuscript write-up.

