## Supplementary Material for "COALIA: a computational model of human EEG for consciousness research"

### 1 Modeling of micro- and macro-circuits: Neural mass model approach

The neural mass model (NMM) represents subsets of excitatory and inhibitory neurons interacting through feedback loops. The physiological properties of each subpopulation are described by two key functions, namely, a “pulse-to-wave” function converting the average pre-synaptic pulse density of afferent action potentials into an average postsynaptic membrane potential, and a “wave-to-pulse” function that converts the average postsynaptic potential into an average pulse density of action potentials. The “pulse-to-wave” function can be expressed as a 2^nd^ order linear filter expressed as $h\left( t \right)=Wwte^{-wt}$, where *W* and *w* are the amplitude and time constant of the average receptor-mediated postsynaptic potential, respectively (Freeman 1992). The pulse-to-wave function $h\left( t \right)$ thereby introduces a 2^nd^ order ordinary differential equation (ODE) of the form: $\ddot{y}\left( t \right)=Wwx\left( t \right)-2w\dot{y}(t)-w^{2}y\left( t \right)$, where $x\left( t \right)$and $y\left( t \right)$ are the input (afferent pulse density) and output (average postsynaptic membrane potential) signals, respectively. The “wave-to-pulse” function is instead modeled by a static nonlinear sigmoid-shaped function $S\left( v \right)={Q_{max}}/{(1+e^{r(v_{0}-v)})}$ that characterizes the saturation and threshold effects taking place at the soma level. The interactions between the different subpopulations are modeled by connectivity constants representing the average number of synaptic contacts between the considered sub-populations. In the following, given a source subpopulation “S” and a target one “T” in NMM of index “*n*”, the connectivity constant projecting from “S” to “T” is denoted$C_{S,T}^{n}$. In the case of direct coupling (“S” and “T” are the same), the notation is reduced to$C_{S}^{n}$. The computational model consisted of a neocortical module containing *N* NMMs, a thalamic module and their interactions.

#### 1.1 ODEs of the neocortical module

The variables$v_{P}^{n}$,$v_{P'}^{n}$, $v_{BC}^{n}$,$v_{SST}^{n}$ and $v_{VIP}^{n}$ are the outputs of the set of ODEs in the neocortical module. They refer to the average postsynaptic membrane potentials at the level of PC, PC’, BC, SST and VIP in the *n*^th^ neural mass model, respectively. $p_{f,c}^{n}\left( t \right)$ refers to the filtered pyramidal noise, so that$\ddot{p}_{f,c}^{n}= A_{c}a_{c}p_{c}^{n}\left( t \right)-2a_{c}\dot{p}_{f,c}^{n}-a_{c}^{2} p_{f,c}^{n}$. The variables$w_{P,X}^{n}$ and$w_{Th,X}^{n}$ refer to the excitatory postsynaptic potentials (EPSP) originating from PC and TC, respectively, and projecting to subpopulation *X* in NMM of index “*n*” (see section 1.3 for details).

$\ddot{v}_{P}^{n}= A_{c}a_{c}S\left( w_{P,P}^{n}+w_{Th,P}^{n}+v_{P'}^{n}-C_{BC,P}^{n}v_{BC}-C_{SST,P}^{n}v_{SST}+p_{f,c}^{n}\left( t \right) \right)-2a_{c}\dot{v}_{P}^{n}-a_{c}^{2} v_{P}^{n}$	(1)

$\ddot{v}_{P'}^{n}= A_{c}a_{c}\left[ C_{P',P}^{n}S\left( C_{P,P'}^{n}v_{P}^{n} \right) \right]-2a_{c}\dot{v}_{P'}^{n}-a_{c}^{2} v_{P'}^{n}$	(2)

$\ddot{v}_{BC}^{n}= G_{c}g_{c}S\left( w_{P,BC}^{n}+w_{Th,BC}^{n}+C_{P,BC}^{n}v_{P}^{n}-C_{SST,BC}^{n}v_{SST}^{n}-{C_{BC}^{n}v}_{BC}^{n} \right)-2g_{c}\dot{v}_{BC}^{n}-g_{c}^{2} v_{BC}^{n}$	(3)

$\ddot{v}_{SST}^{n}= B_{c}b_{c}S\left( w_{P,SST}^{n}+w_{Th,SST}^{n}-C_{VIP,SST}^{n}v_{VIP}^{n} \right)-2b_{c}\dot{v}_{SST}^{n}-b_{c}^{2} v_{SST}^{n}$	(4)

$\ddot{v}_{VIP}^{n}= D_{c}d_{c}S\left( w_{P,VIP}^{n}+w_{Th,VIP}^{n}-C_{SST,VIP}^{n}v_{SST}^{n} \right)-2d_{c}\dot{v}_{VIP}^{n}-d_{c}^{2} v_{VIP}^{n}$	(5)

#### 1.2 ODEs of the thalamic module

The variables$v_{Th}$, $v_{{TRN}_{1}}$and $v_{{TRN}_{2}}$ are the outputs of the set of ODEs in the thalamic module. They refer to the average postsynaptic membrane potentials at the level of TC, TRN_1_ and TRN_2_ cells, respectively. $p_{f,Th}^{n}\left( t \right)$ refers to the filtered thalamic noise, so that$\ddot{p}_{f,Th}^{n}= A_{Th}a_{Th}p_{Th}^{n}\left( t \right)-2a_{Th}\dot{p}_{f,Th}^{n}-a_{c}^{2} p_{f,Th}^{n}$. The variable $w_{P,X}$ refers to the EPSP originating from PC and projecting to subpopulation *X* in the thalamic module (see section 1.3 for details).

$\ddot{v}_{Th}= A_{Th}a_{Th}S\left( w_{P,Th}+C_{Th}v_{Th}-C_{{TRN}_{1},Th}v_{{TRN}_{1}}-C_{{TRN}_{2},Th}v_{{TRN}_{2}}+p_{f,Th}\left( t \right) \right)-2a_{Th}\dot{v}_{Th}-a_{Th}^{2} v_{Th}$	(6)

$\ddot{v}_{{TRN}_{1}}= G_{Th}g_{Th}S\left( w_{P,{TRN}_{1}}+C_{Th,{TRN}_{1}}v_{Th} \right)-2g_{Th}\dot{v}_{{TRN}_{1}}-g_{Th}^{2} v_{{TRN}_{1}}$	(7)

$\ddot{v}_{{TRN}_{2}}= B_{Th}b_{Th}S\left( w_{P,{TRN}_{2}}+C_{Th,{TRN}_{2}}v_{Th} \right)-2b_{Th}\dot{v}_{{TRN}_{2}}-b_{Th}^{2} v_{{TRN}_{2}}$	(8)

#### 1.3 ODEs of large scale cortico- and thalamo-cortical connectivity

In the model, two types of large-scale connectivity were considered, namely, the cortico-cortical connectivity relating neocortical NMMs and implemented by equations (9)-(12), and the thalamo-cortical connectivity describing the thalamo-cortical loop and implemented by equations (13)-(19).

$\ddot{w}_{P,P}^{n}= A_{c}a_{c}S\left( \sum_{i=1,i\neq n}^{N} K_{P,P}^{i,n}v_{P}^{i} \right)-2a_{c}\dot{w}_{P,P}^{n}-a_{c}^{2} w_{P,P}^{n}$	(9)

$\ddot{w}_{P,BC}^{n}= A_{c}a_{c}S\left( \sum_{i=1,i\neq n}^{N} K_{P,BC}^{i,n}v_{P}^{i} \right)-2a_{c}\dot{w}_{P,BC}^{n}-a_{c}^{2} w_{P,BC}^{n}$	(10)

$\ddot{w}_{P,SST}^{n}= A_{c}a_{c}S\left( \sum_{i=1,i\neq n}^{N} K_{P,SST}^{i,n}v_{P}^{i} \right)-2a_{c}\dot{w}_{P,SST}^{n}-a_{c}^{2} w_{P,SST}^{n}$	(11)

$\ddot{w}_{P,VIP}^{n}= A_{c}a_{c}S\left( \sum_{i=1,i\neq n}^{N} K_{P,VIP}^{i,n}v_{P}^{i} \right)-2a_{c}\dot{w}_{P,VIP}^{n}-a_{c}^{2} w_{P,VIP}^{n}$	(12)

$\ddot{w}_{Th,P}^{n}= A_{c}a_{c}S\left( K_{Th,P}^{n}v_{Th} \right)-2a_{c}\dot{w}_{Th,P}^{n}-a_{c}^{2} w_{Th,P}^{n}$	(13)

$\ddot{w}_{Th,BC}^{n}= A_{c}a_{c}S\left( K_{Th,BC}^{n}v_{Th} \right)-2a_{c}\dot{w}_{Th,BC}^{n}-a_{c}^{2} w_{Th,BC}^{n}$	(14)

$\ddot{w}_{Th,SST}^{n}= A_{c}a_{c}S\left( K_{Th,SST}^{n}v_{Th} \right)-2a_{c}\dot{w}_{Th,SST}^{n}-a_{c}^{2} w_{Th,SST}^{n}$	(15)

$\ddot{w}_{Th,VIP}^{n}= A_{c}a_{c}S\left( K_{Th,VIP}^{n}v_{Th} \right)-2a_{c}\dot{w}_{Th,VIP}^{n}-a_{c}^{2} w_{Th,VIP}^{n}$	(16)

$\ddot{w}_{P,Th}= A_{Th}a_{Th}S\left( \sum_{i=1}^{N} K_{P,Th}^{i}v_{P}^{i} \right)-2a_{Th}\dot{w}_{P,Th}-a_{Th}^{2} w_{P,Th}$	(17)

$\ddot{w}_{P,{TRN}_{1}}= A_{Th}a_{Th}S\left( \sum_{i=1}^{N} K_{P,{TRN}_{1}}^{i}v_{P}^{i} \right)-2a_{Th}\dot{w}_{P,{TRN}_{1}}-a_{Th}^{2} w_{P,{TRN}_{1}}$	(18)

$\ddot{w}_{P,{TRN}_{2}}= A_{Th}a_{Th}S\left( \sum_{i=1}^{N} K_{P,{TRN}_{2}}^{i}v_{P}^{i} \right)-2a_{Th}\dot{w}_{P,{TRN}_{2}}-a_{Th}^{2} w_{P,{TRN}_{2}}$	(19)

**Table 1.** Cortico- and thalamo-cortical models: parameters, values and interpretation

| *Cortical module* | | |
| --- | --- | --- |
| Parameter | **Value** | **Interpretation** |
| $A_{c}$ | 4 mV | Amplitude of the cortical average EPSP |
| $B_{c}$ | 30 mV | Amplitude of the cortical average IPSP (GABA*_A_*_,_ *_slow_* mediated currents) |
| $G_{c}$ | 20 mV | Amplitude of the cortical average IPSP (GABA_A, fast_ mediated currents) |
| $D_{c}$ | 10 mV | Amplitude of the cortical average IPSP (GABA_A, slow_ mediated currents) |
| ${1/a}_{c}$ | 1/100 s | Time constant of cortical glutamate-mediated synaptic transmission |
| ${1/b}_{c}$ | 1/30 s | Time constant of cortical GABA-mediated synaptic transmission (GABA*_A, slow_* receptors) |
| ${1/g}_{c}$ | 1/150 s | Time constant of cortical GABA-mediated synaptic transmission (GABA*_A, fast_* receptors) |
| ${1/d}_{c}$ | 1/20 s | Time constant of cortical GABA-mediated synaptic transmission (GABA*_A, slow_* receptors) |
| $\mu_{c}$, $\sigma_{c}$ | 1s^-1^, 1 s^-1^ | Mean and standard deviation of nonspecific cortical input |
| $c$ | 0.5 m.s^-1^ | Average conduction velocity of action potentials |
| $C_{P,P'}^{n}$ | 135 | Collateral excitation connectivity constant of *n*^th^ cortical population |
| $C_{P',P}^{n}$ | 100 | Collateral excitation connectivity constant of *n*^th^ cortical population |
| $C_{BC,P}^{n}$ | 50 | BC to PC connectivity constant of *n*^th^ cortical population |
| $C_{SST,P}^{n}$ | 20 | SST to PC connectivity constant of *n*^th^ cortical population |
| $C_{P,BC}^{n}$ | 50 | PC to BC connectivity constant of *n*^th^ cortical population |
| $C_{P,SST}^{n}$ | 50 | PC to SST connectivity constant of *n*^th^ cortical population |
| $C_{SST,BC}^{n}$ | 13.5 | SST to BC connectivity constant of *n*^th^ cortical population |
| $C_{SST,VIP}^{n}$ | 20 | SST to VIP connectivity constant of *n*^th^ cortical population |
| $C_{VIP,SST}^{n}$ | 20 | VIP to SST connectivity constant of *n*^th^ cortical population |
| $C_{BC}^{n}$ | 20 | BC to BC connectivity constant of *n*^th^ cortical population |
| *Thalamic module* | | |
| Parameter | **Value** | **Interpretation** |
| $A_{Th}$ | 4.1 mV | Amplitude of the thalamic average EPSP |
| $B_{Th}$ | 5 mV | Amplitude of the thalamic average IPSP (GABA*_A, slow_* and GABA*_B_* receptors) |
| $G_{Th}$ | 20 mV | Amplitude of the thalamic average IPSP (GABA*_A, fast_* receptors) |
| ${1/a}_{Th}$ | 1/100 s | Time constant of thalamic glutamate-mediated synaptic transmission |
| ${1/b}_{Th}$ | 1/20 s | Time constant of thalamic GABA-mediated synaptic transmission (GABA*_A, slow_* and GABA*_B_* receptors) |
| ${1/g}_{Th}$ | 1/150 s | Time constant of thalamic GABA-mediated synaptic transmission (GABA*_A, fast_* receptors) |
| $\mu_{Th}$, $\sigma_{Th}$ | 120 s^-1^, 30 s^-1^ | Mean and standard deviation of nonspecific subcortical input |
| $C_{Th,{TRN}_{1}}$ | 20 | TC to TRN_1_ connectivity constant |
| $C_{Th,{TRN}_{2}}$ | 20 | *TC* to TRN_2_ connectivity constant |
| $C_{{TRN}_{1},Th}$ | 10 | TRN_1_ to TC connectivity constant |
| $C_{{TRN}_{2},Th}$ | 10 | TRN_2_ to TC connectivity constant |
| $C_{Th}$ | 30 | Collateral excitation connectivity constant |
| *Cortico- and thalamo-cortical connections* | | |
| Parameter | **Value** | **Interpretation** |
| $K_{P,P}^{i,n}$ |  | PC to PC connectivity constant from *i*^th^ to *n*^th^ cortical populations |
| $K_{P,BC}^{i,n}$ |  | PC to BC connectivity constant from *i*^th^ to *n*^th^ cortical populations |
| $K_{P,SST}^{i,n}$ |  | PC to SST connectivity constant from *i*^th^ to *n*^th^ cortical populations |
| $K_{P,VIP}^{i,n}$ |  | PC to VIP connectivity constant from *i*^th^ to *n*^th^ cortical populations |
| $K_{Th,P}^{n}$ |  | TC to PC connectivity constant of *n*^th^ cortical population |
| $K_{P,Th}^{n}$ |  | PC of *n*^th^ cortical population to TC connectivity constant |
| $K_{Th,BC}^{n}$ |  | TC to BC connectivity constant of *n*^th^ cortical population |
| $K_{Th,VIP}^{n}$ |  | TC to VIP connectivity constant of *n*^th^ cortical population |
| $K_{Th,SST}^{n}$ |  | TC to SST connectivity constant of *n*^th^ cortical population |
| $K_{P,{TRN}_{1}}^{n}$ |  | PC of *n*^th^ cortical population to TRN1 connectivity constant |
| $K_{P,{TRN}_{2}}^{n}$ |  | PC of *n*^th^ cortical population to TRN2 connectivity constant |
| $v_{0}$, $e_{0}$, $r$ | 6 mV, 2.5 s^-1^, 0.56 mV^-1^ | Parameters of the nonlinear sigmoid function (transforming the average membrane potential to an average density of action potentials) |

### 2 Large-scale cortico-cortical and thalamocortical connectivity

$\boldsymbol{K}_{\boldsymbol{EXC}}$	$\boldsymbol{K}_{\boldsymbol{BC}}$

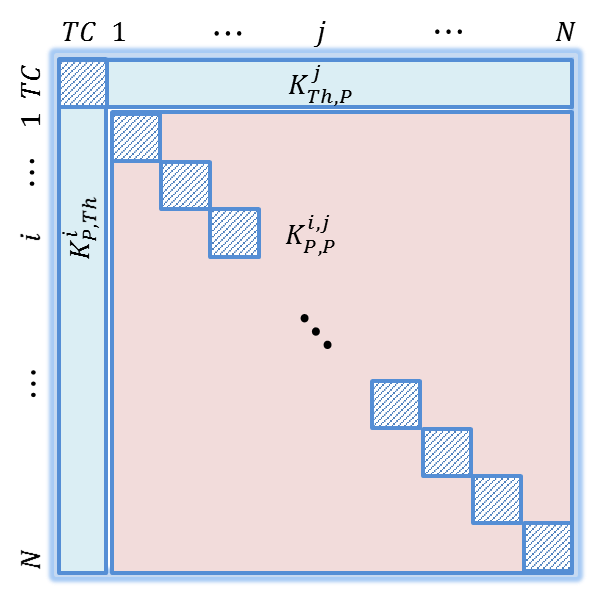

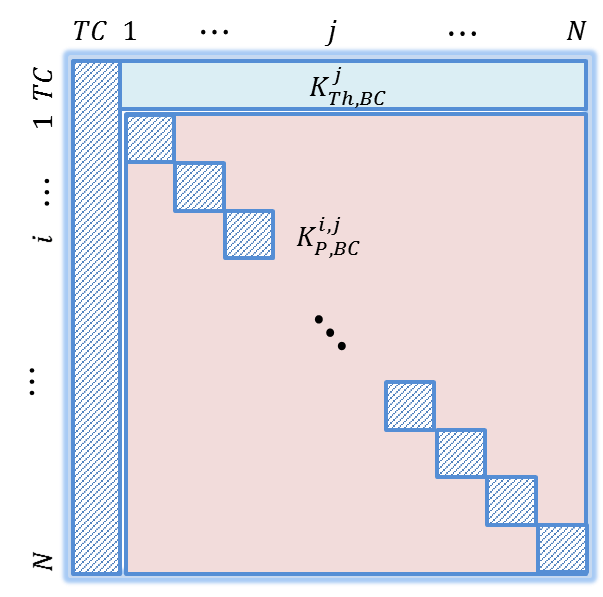

$\boldsymbol{K}_{\boldsymbol{SST}}$	$\boldsymbol{K}_{\boldsymbol{VIP}}$

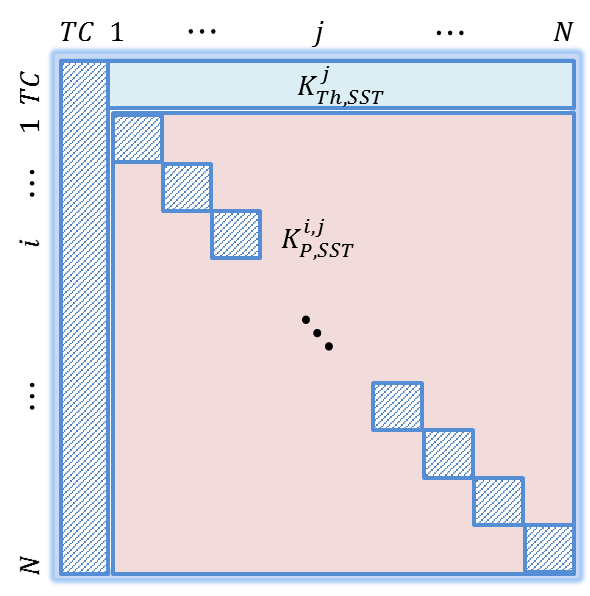

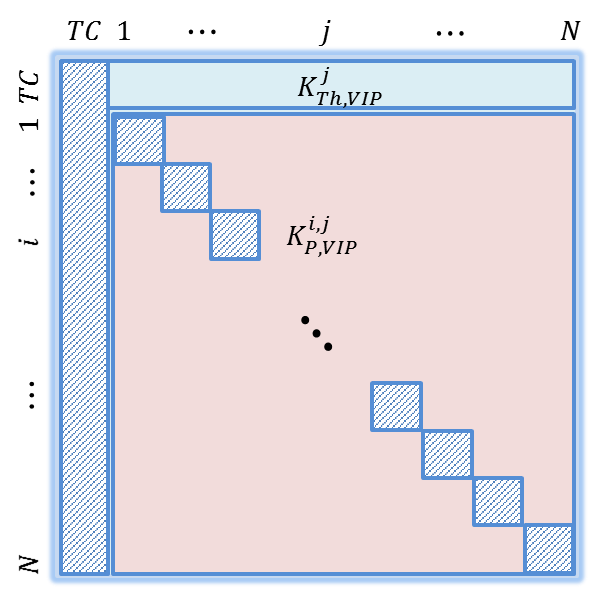

$\boldsymbol{K}_{\boldsymbol{TRN}_{\boldsymbol{1}}}$ $\boldsymbol{K}_{\boldsymbol{TRN}_{\boldsymbol{2}}}$

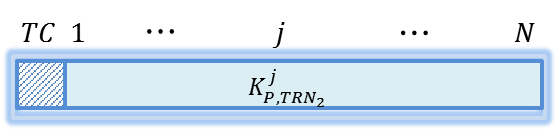

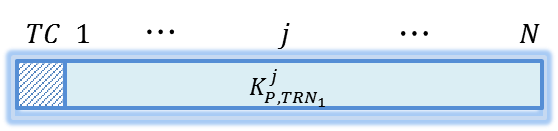

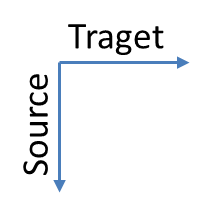

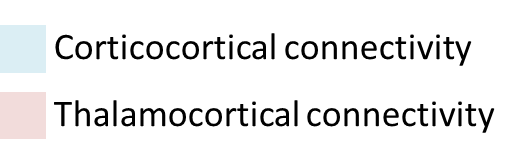

Figure 1. Large-scale connectivity matrices. Given one thalamic population (NMM “0”) and *N* cortical NMMs, inter-population connectivity constants were arranged in six matrices: 1) four two-dimensional matrices of size$\boldsymbol{(N+1)\times(N+1)}$, namely,$\boldsymbol{K}$,$\boldsymbol{K}_{\boldsymbol{BC}}$, $\boldsymbol{K}_{\boldsymbol{SST}}$ and $\boldsymbol{K}_{\boldsymbol{VIP}}$; and 2) two one-dimensional matrices of length $\boldsymbol{(N+1)}$, $\boldsymbol{K}_{\boldsymbol{TRN}_{\boldsymbol{1}}}$ and $\boldsymbol{K}_{\boldsymbol{TRN}_{\boldsymbol{2}}}$. Hashed components refer to irrelevant connections, such as the diagonal elements (when “*i*” = “*j*”) since they do not carry inter-population connections, and the first columns in$\boldsymbol{K}_{\boldsymbol{BC}}$, $\boldsymbol{K}_{\boldsymbol{SST}}$ and $\boldsymbol{K}_{\boldsymbol{VIP}}$since NMM “0” is the thalamic population. Consequently, these connections are automatically set to 0.

#### 2.1 Connectivity matrices used in the toy model

| $\boldsymbol{K}_{\boldsymbol{EXC}}$  $\boldsymbol{K}_{\boldsymbol{BC}}$  $\boldsymbol{K}_{\boldsymbol{SST}}$  $\boldsymbol{K}_{\boldsymbol{VIP}}$  $\boldsymbol{K}_{\boldsymbol{TRN}_{\boldsymbol{1}}}$  $\boldsymbol{K}_{\boldsymbol{TRN}_{\boldsymbol{2}}}$ | **Deep sleep (SWS) Wakefulness (BKG)**  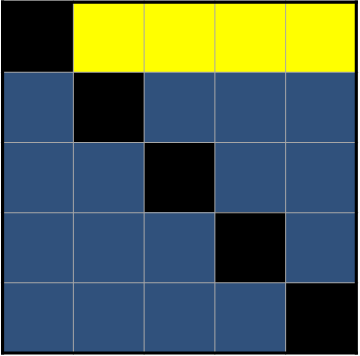 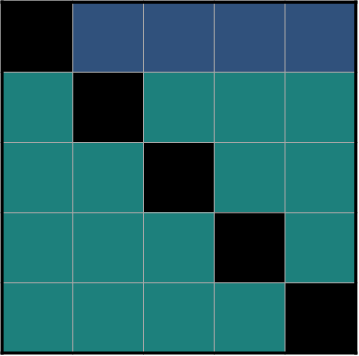  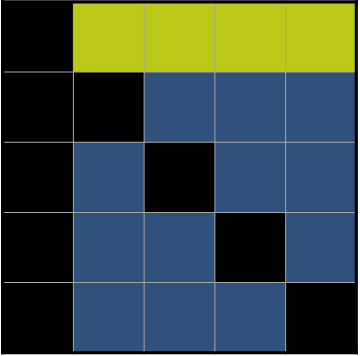 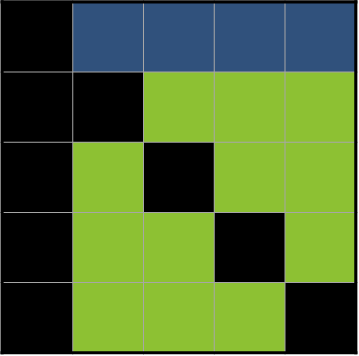  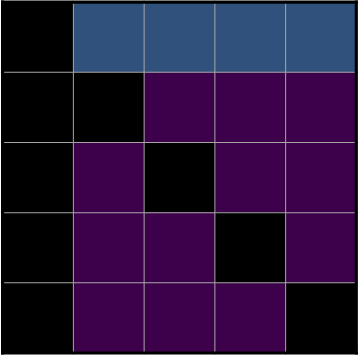 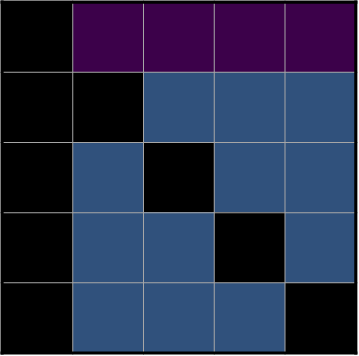  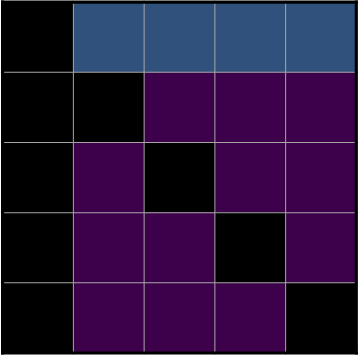 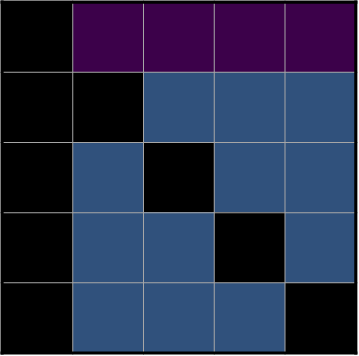  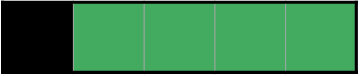 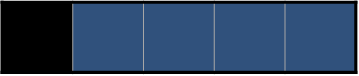  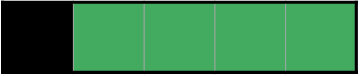 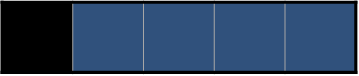 | 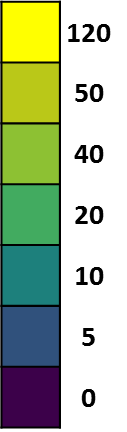 |
| --- | --- | --- |

Figure 2. Connectivity matrices used in the toy model to generate slow wave sleep (SWS) and background activity (BKG).

#### 2.2 Connectivity matrices used in the whole brain model

| $\mathcal{K}_{\boldsymbol{EXC}}$  $\mathcal{K}_{\boldsymbol{BC}}$  $\mathcal{K}_{\boldsymbol{SST}}$  $\mathcal{K}_{\boldsymbol{VIP}}$  $\mathcal{K}_{\boldsymbol{TRN}_{\boldsymbol{1}}}$  $\mathcal{K}_{\boldsymbol{TRN}_{\boldsymbol{2}}}$ | **Deep sleep (SWS) Wakefulness (BKG)**  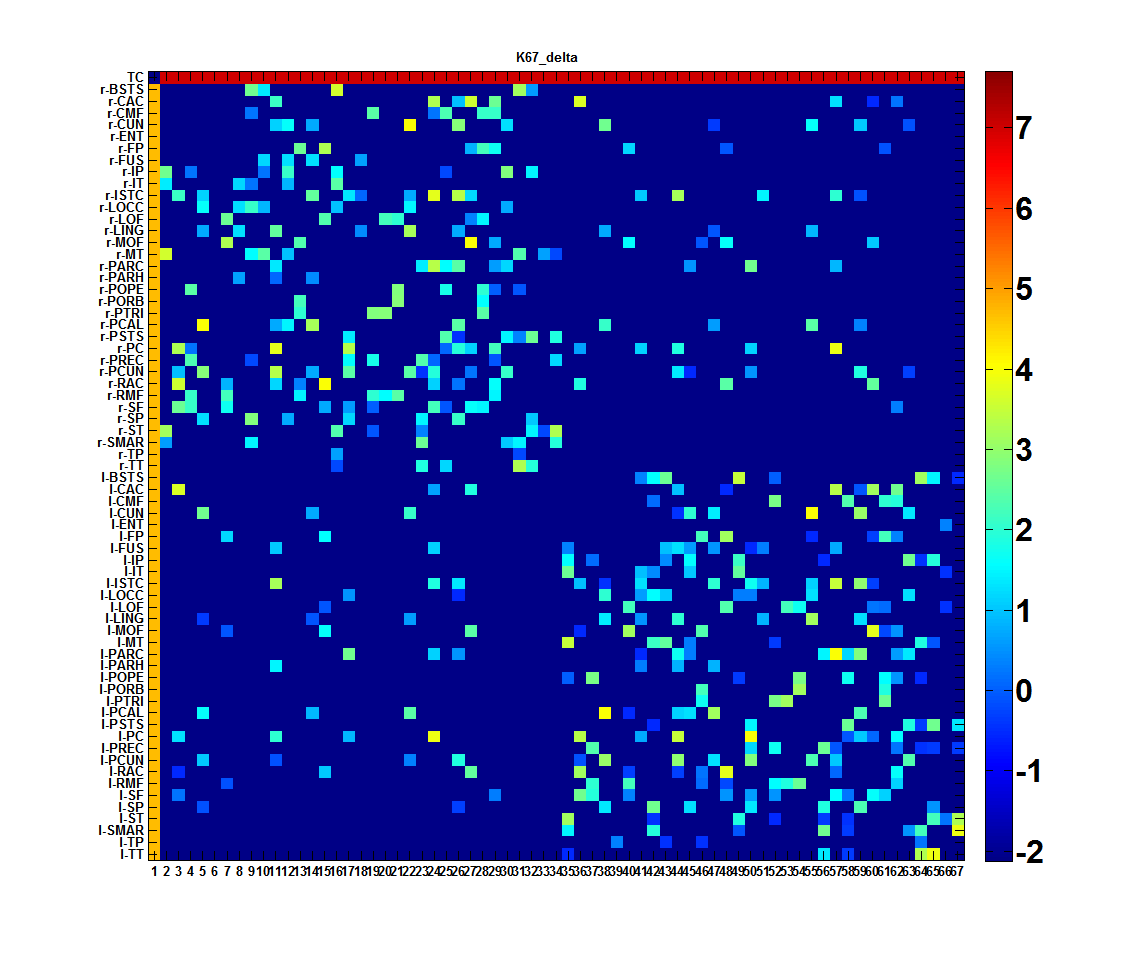 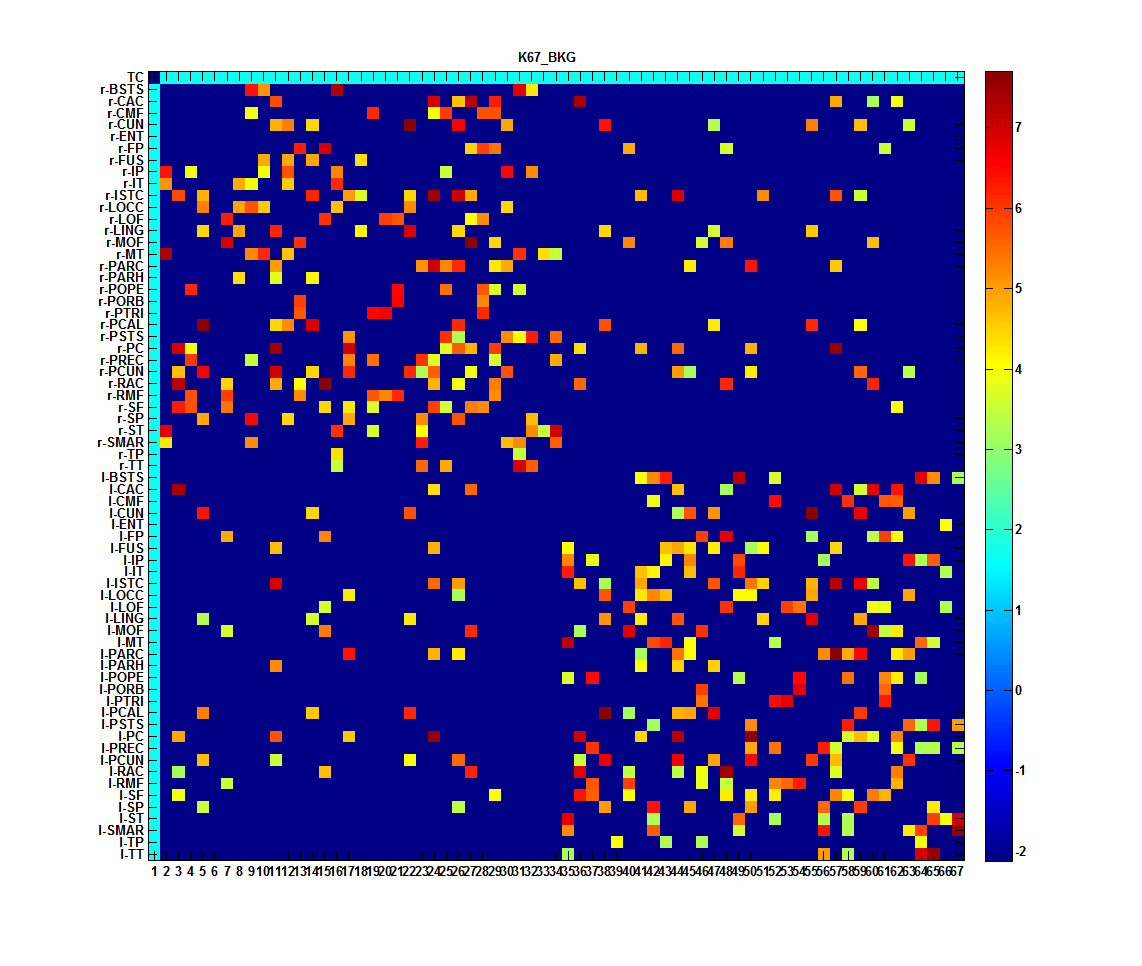  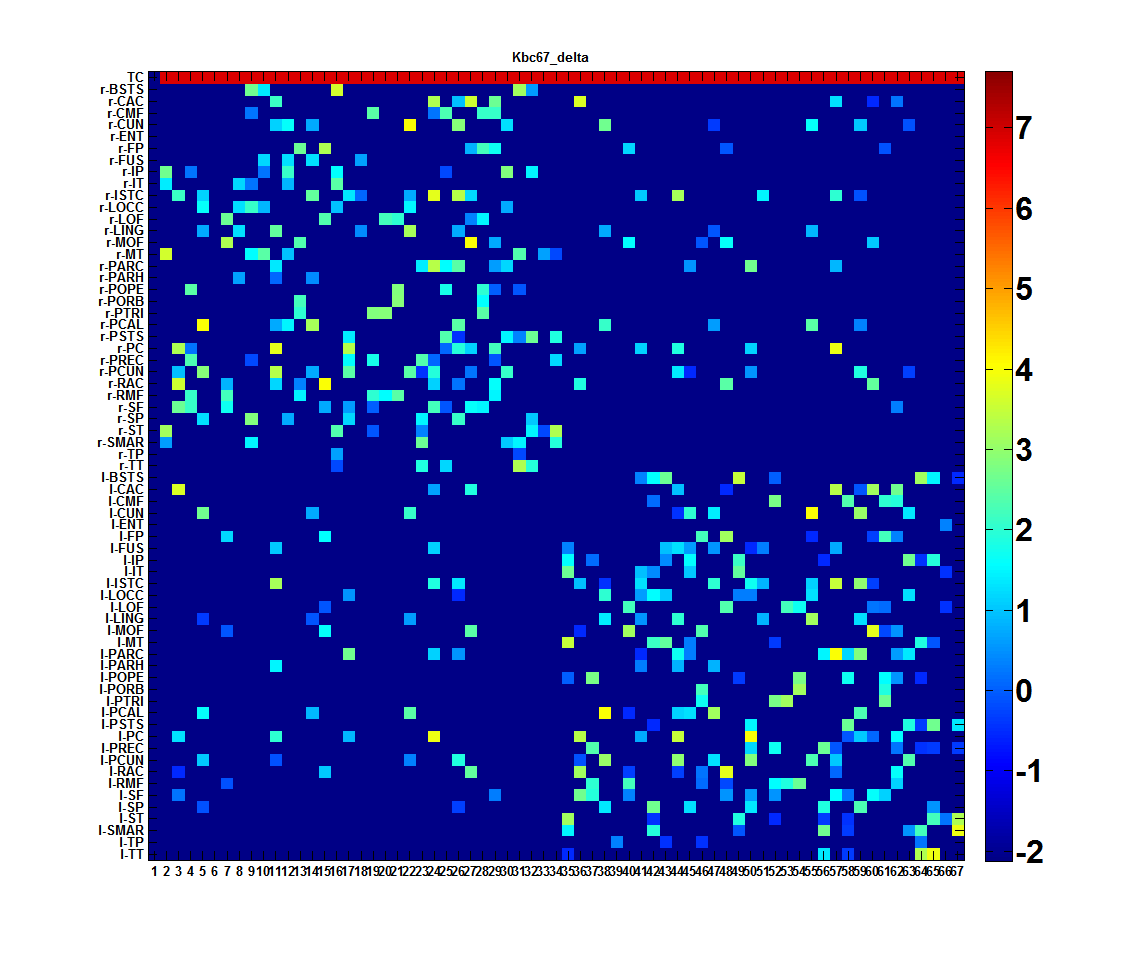 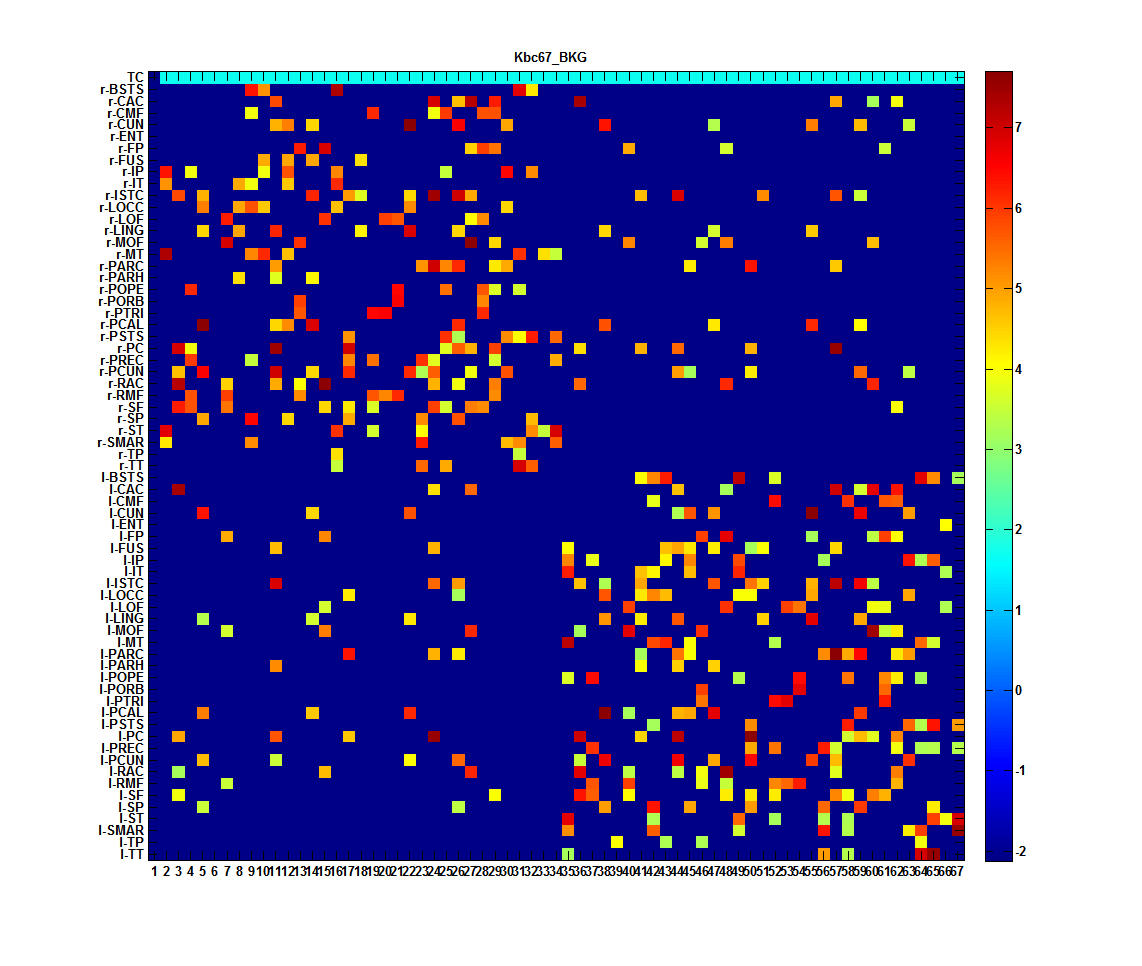  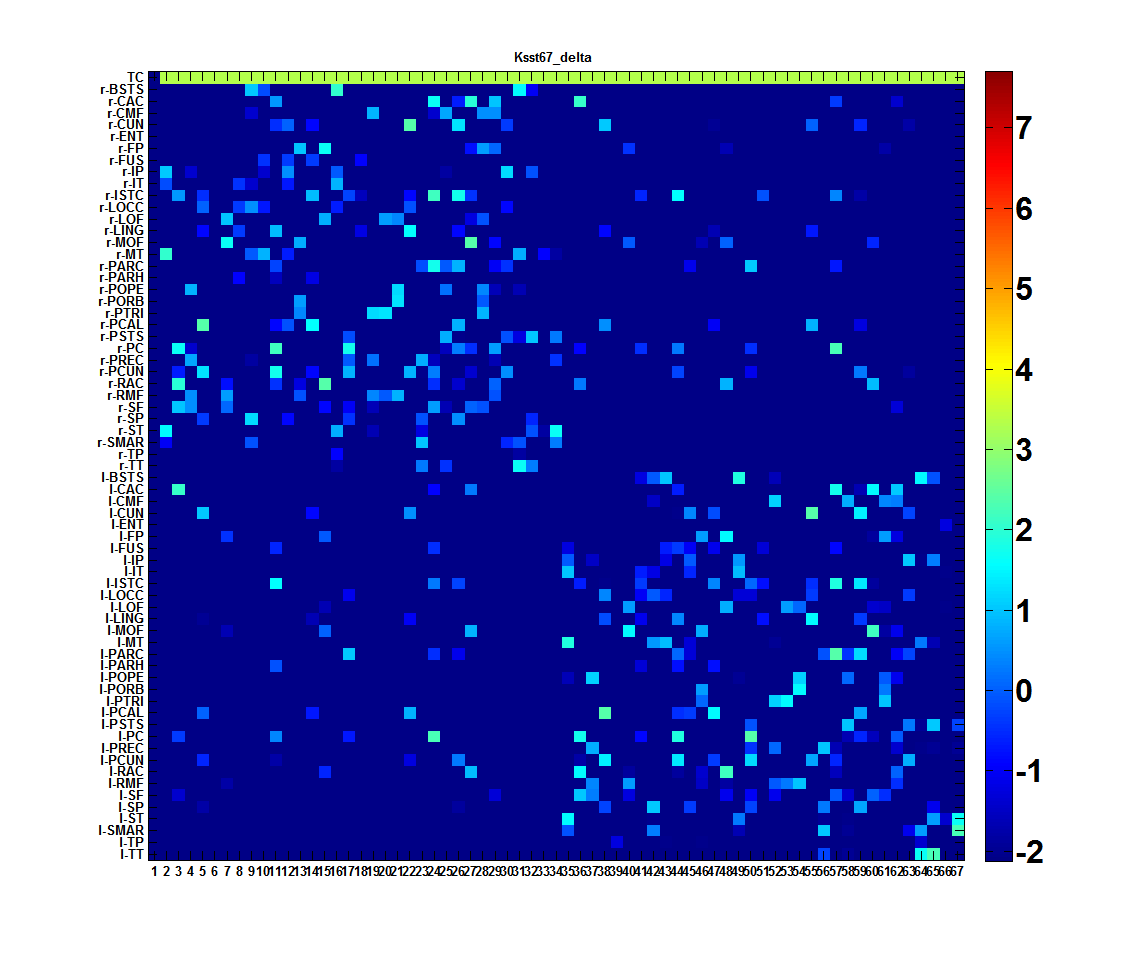 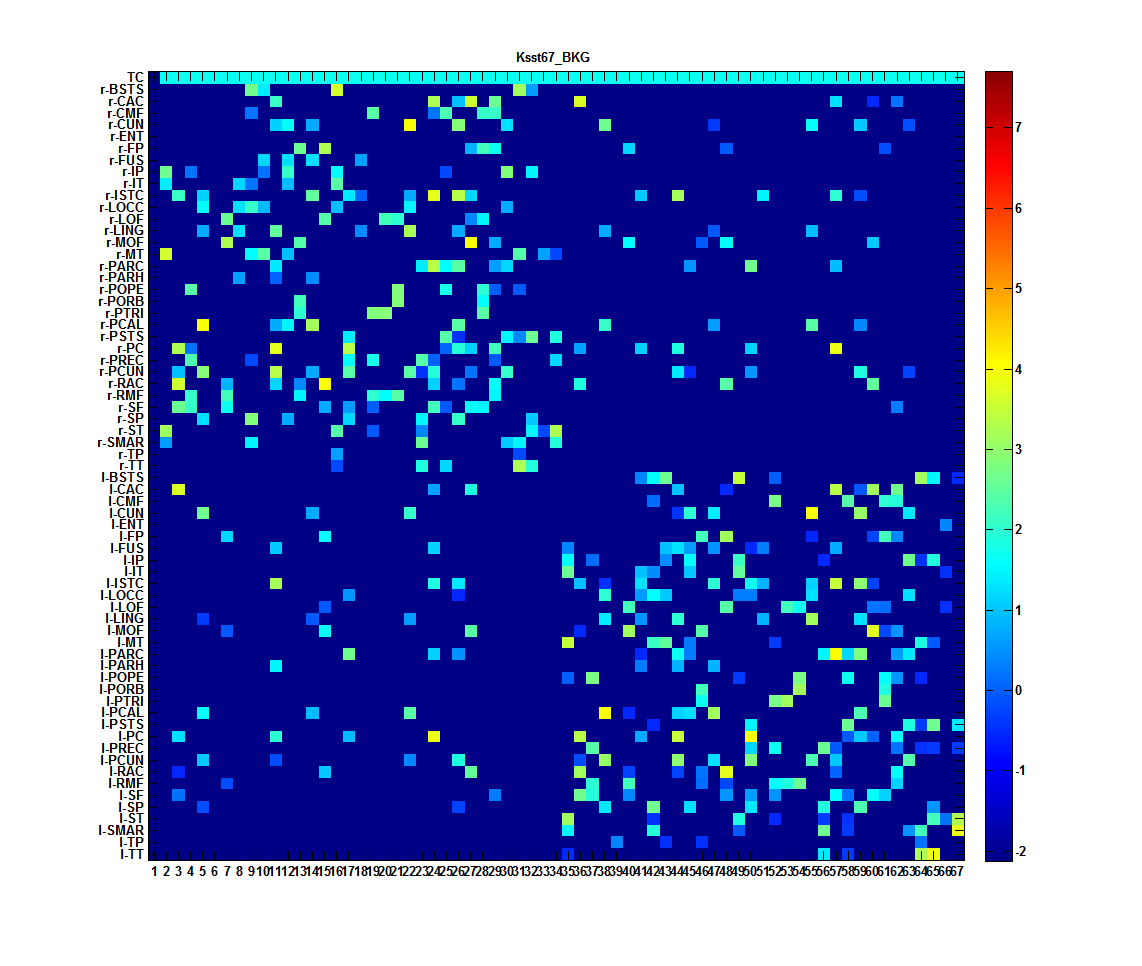  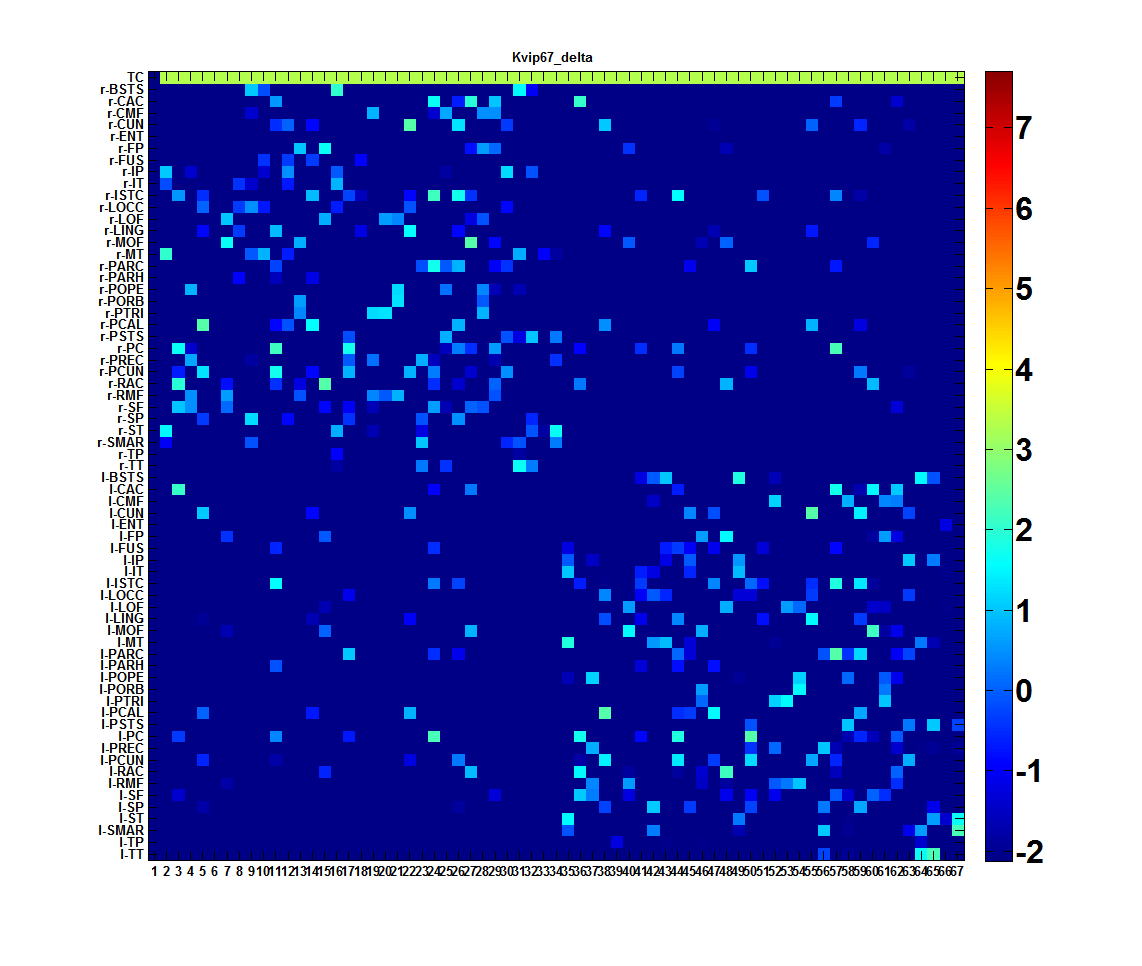 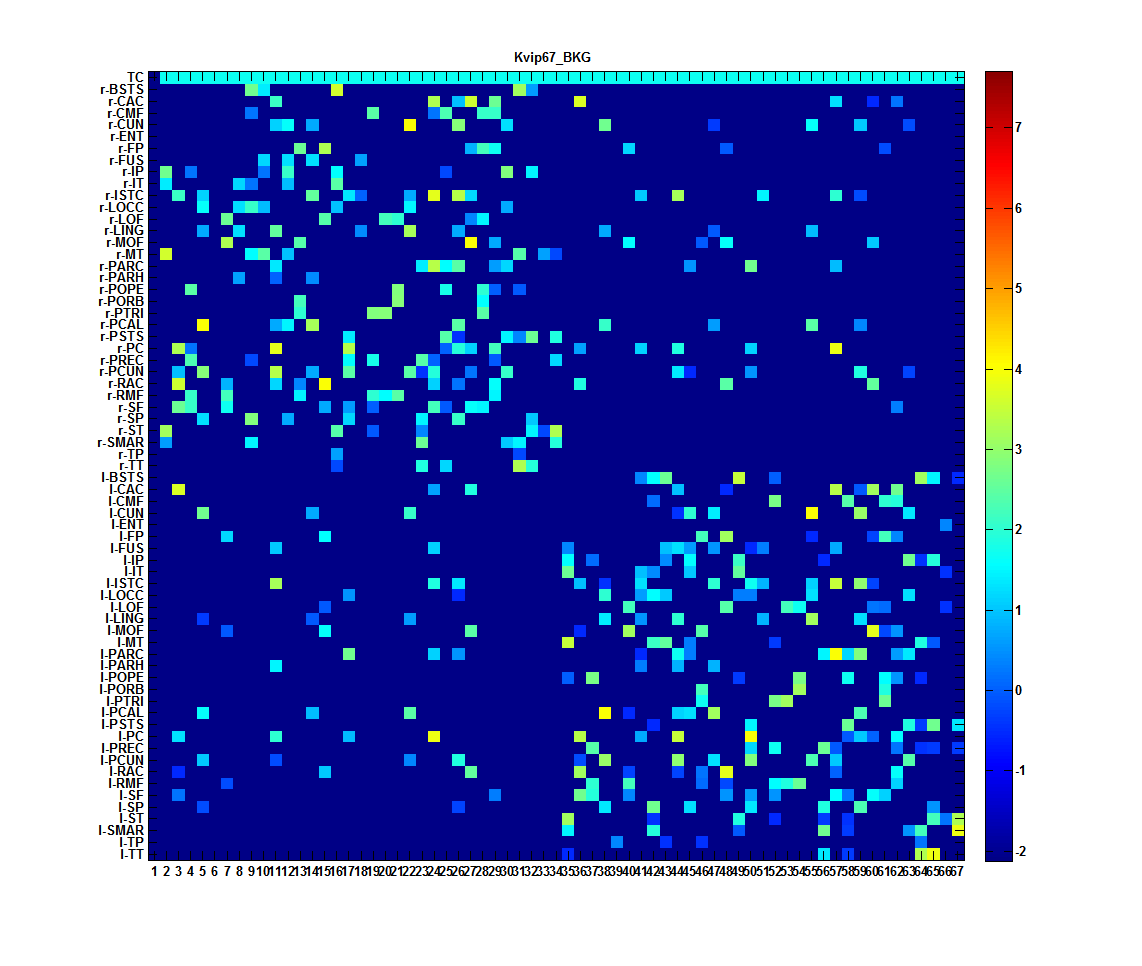  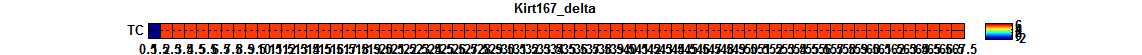 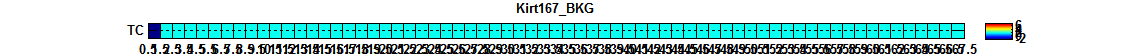  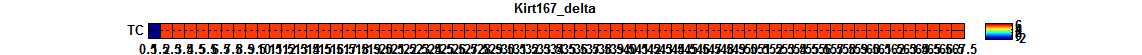 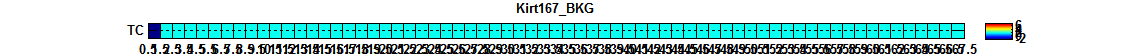 | 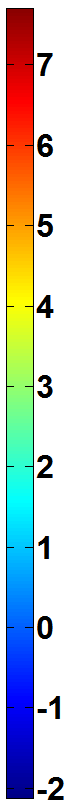 |
| --- | --- | --- |

Figure 3. Connectivity matrices used in the whole brain model to generate slow wave sleep (SWS) and background activity (BKG). The logarithmic amplitude of matrices are displayed in order to highlight the difference between the two states (whence the negative values of connectivity).
